# Predicting the Key Regulators of Cell Identity in Human Adult Pancreas

**DOI:** 10.1101/2020.09.23.310094

**Authors:** Lotte Vanheer, Federica Fantuzzi, San Kit To, Andrea Alex Schiavo, Matthias Van Haele, Tine Haesen, Xiaoyan Yi, Adrian Janiszewski, Joel Chappell, Adrien Rihoux, Toshiaki Sawatani, Tania Roskams, Francois Pattou, Julie Kerr-Conte, Miriam Cnop, Vincent Pasque

## Abstract

Cellular identity during development is under the control of transcription factors that form gene regulatory networks. However, the transcription factors and gene regulatory networks underlying cellular identity in the human adult pancreas remain largely unexplored. Here, we integrate multiple single-cell RNA-sequencing datasets of the human adult pancreas, totaling 7393 cells, and comprehensively reconstruct gene regulatory networks. We show that a network of 142 transcription factors forms distinct regulatory modules that characterize pancreatic cell types. We present evidence that our approach identifies regulators of cell identity in the human adult pancreas. We predict that HEYL, BHLHE41 and JUND are active in acinar, beta and alpha cells, respectively, and show that these proteins are present in the human adult pancreas as well as in human induced pluripotent stem cell (hiPSC)-derived islet cells. Using single-cell transcriptomics, we found that JUND represses beta cell genes in hiPSC-alpha cells. Both BHLHE41 and JUND depletion seemed to increase the number of sc-enterochromaffin cells in hiPSC-derived islets. The comprehensive gene regulatory network atlas can be explored interactively online. We anticipate our analysis to be the starting point for a more sophisticated dissection of how transcription factors regulate cell identity in the human adult pancreas. Furthermore, given that transcription factors are major regulators of embryo development and are often perturbed in diseases, a comprehensive understanding of how transcription factors work will be relevant in development and disease.

**HIGHLIGHTS:**

- Reconstruction of gene regulatory networks for human adult pancreatic cell types
- An interactive resource to explore and visualize gene expression and regulatory states
- Prediction of putative transcription factors that drive pancreatic cell identity
- BHLHE41 depletion in primary islets induces apoptosis

## INTRODUCTION

A fundamental question in biology is how a single genome gives rise to the great diversity of cell types that make up organs and tissues. A key goal is to map all cell types of developing and mature organs such as the pancreas, an essential organ at the basis of multiple human disorders including diabetes and cancer (Kahn, Cooper, and Del Prato 2014; Kamisawa et al. 2016; Han et al. 2020). Single-cell RNA-sequencing (scRNA-seq) provides a powerful tool to resolve cellular heterogeneity, identify cell types and capture high-resolution snapshots of gene expression in individual cells (Grün et al. 2016). With the advent of single-cell transcriptomics, great progress has been made toward the creation of a reference cell atlas of the pancreas (Baron et al. 2016; Segerstolpe et al. 2016; Y. J. Wang et al. 2016; Xin et al. 2016; Enge et al. 2017; Lawlor et al. 2017; Augsornworawat et al. 2020; Han et al. 2020; Augsornworawat and Millman 2020; S. He et al. 2020; Fasolino et al. 2022). Work from several groups provided cellular atlases of the pancreas during mouse development (Stanescu et al. 2017; Byrnes et al. 2018; Scavuzzo et al. 2018; Duvall et al. 2022), in adult mice (Muraro et al. 2016) and in human fetal (Cao et al. 2020; la O Sean et al. 2022; Gonçalves et al. 2021; Y. Xu et al. 2022; Olaniru et al. 2022; Lin et al. 2021) and adult pancreas (Liu et al. 2014; Baron et al. 2016; Muraro et al. 2016; Segerstolpe et al. 2016; Y. J. Wang et al. 2016; Xin et al. 2016; Lawlor et al. 2017; Enge et al. 2017; Han et al. 2020; Kang et al. 2022; Wigger et al. 2021). Efforts have also been made to map cellular identity during pancreas development starting from human pluripotent stem cells (Gonçalves et al. 2021; Rezania et al. 2014; Hrvatin et al. 2014; Z. Zhu et al. 2016; Augsornworawat et al. 2020; Hogrebe et al. 2021; Peterson et al. 2020; Veres et al. 2019; Balboa et al. 2022; Krentz et al. 2018; Kang et al. 2022; Augsornworawat et al. 2022; Budnik et al. 2022). Taken together, these studies provide an opportunity to better understand the establishment and maintenance of cellular identity among different pancreatic cell types.

Work over the past decades indicated that cellular identity is established by combinations of transcription factors (TFs) that recognize and interact with cis-regulatory elements in the genome (Fiers et al. 2018). TFs, together with chromatin modifiers, deploy gene expression programs. A small number of core TFs are thought to be sufficient for the establishment and maintenance of gene expression programs that define cellular identity during and after development (Ohno 1979). Studies conducted in both mouse and human have successfully identified TFs that are pivotal for the acquisition and maintenance of pancreatic cell fates (Dassaye, Naidoo, and Cerf 2016). These include PDX1 (Zhou et al. 2008; Shih et al. 2015), MAFA (Nishimura, Takahashi, and Yasuda 2015), NGN3 (Gradwohl et al. 2000), NKX2.2 (Sussel et al. 1998), PAX4 (Sosa-Pineda et al. 1997), NKX6.1, NEUROD1 (Mastracci et al. 2013), ARX (Collombat et al. 2003), MAFB (Artner et al. 2006), RFX6 (Smith et al. 2010; Piccand et al. 2014), GATA4 (Ketola et al. 2004), FOXA2 (Lee et al. 2019) and SOX9 (Shroff et al. 2014; Shih et al. 2015). Conditional deletion of TFs such as FOXA2 and PDX1 in adult beta cells results in the loss of cellular identity and function (Sund et al. 2001; Gao et al. 2014). Robust genetic evidence for the role of these TFs in establishing human pancreatic cell identity is provided by the identification of TF loss-of-function mutations that cause pancreatic agenesis (Stanescu et al. 2017; Sellick et al. 2004; Allen et al. 2011; Shaw-Smith et al. 2014; De Franco et al. 2019) or neonatal or young-onset diabetes (Senée et al. 2006; Solomon et al. 2009; Rubio-Cabezas et al. 2010; Shaw-Smith et al. 2014; Bonnefond et al. 2013; Flanagan et al. 2014). In addition, TF overexpression can reprogram somatic cells to adopt alternative identities (Takahashi and Yamanaka 2006; Vierbuchen et al. 2010; Lima et al. 2016). For example, the induced expression of *Ngn3, Pdx1*, and *Mafa* was shown to reprogram mouse alpha cells into beta-like cells *in vivo* (*Zhou et al. 2008*). However, how key TFs underlie the maintenance of cellular identity in the human pancreas remains incompletely understood.

Multiple approaches to reconstruct gene regulatory networks (GRNs) from bulk and single-cell omics data have been developed (Ghazanfar et al. 2016; Fiers et al. 2018; H. Matsumoto et al. 2017; Lim et al. 2016). In particular, it is now possible to combine single-cell transcriptomic data with either cis-regulatory information (Kamimoto, Hoffmann, and Morris 2020; Janky et al. 2014; Aibar et al., 2017; Sande et al. 2020) or chromatin accessibility (Kamimoto, Hoffmann, and Morris 2020; González-Blas et al. 2022) to infer GRNs. Because TFs recognize DNA motifs in the genome, one can measure if inferred target genes are expressed within single cells, and therefore quantify the activity of TFs. Such approaches have revealed the regulatory programs in distinct systems including the Drosophila brain (Davie et al. 2018), cancer (Wouters et al. 2020), during early mouse embryonic (Peng et al. 2020) and pancreas development (H. Zhu et al. 2022), reprogramming to induced pluripotency (Talon et al. 2021), in a mouse cell atlas (Suo et al. 2018) and a human cell atlas (Han et al. 2020).

Analysis of GRNs in the human adult pancreas has identified distinct endocrine and exocrine regulatory states with multiple stable cell states for alpha, beta and ductal cells (Kumar and Vinod 2019). No change in GRN activity of alpha and beta cells was reported in type 2 diabetes or related to body mass index (BMI) (Kumar and Vinod 2019). Previous data show that type 2 diabetic (Faerch, Hulman, and Solomon 2015; Dybala and Hara 2019; Zaharia et al. 2019; Dennis et al. 2019) and non-diabetic human islet preparations vary greatly depending on age (Enge et al. 2017; Westacott et al. 2017) and BMI (Henquin 2018) warranting the exploration of GRNs across multiple integrated datasets. Hence, it remains unclear whether previous GRN findings can be extrapolated to a broader, highly heterogeneous population comprising non-diabetic and type 2 diabetic donors. The development of integration methods provides an opportunity to analyze multiple scRNA-seq studies from multiple laboratories and patients (Butler et al. 2018; Luecken et al., 2022). Additional knowledge on how GRNs maintain cellular identity in the human adult pancreas may further the understanding of disease states and improve ongoing efforts to convert human induced pluripotent stem cells (hiPSCs) into functional, mature beta cells for diabetes treatment.

Here, we build an integrated human pancreas gene regulatory atlas. In this resource, we use single-cell transcriptomes of the human adult pancreas, taking advantage of integration strategies and computational tools to reconstruct GRNs. Our analysis identifies the GRN landscape and candidate regulators that are critical for cellular identity in the human adult pancreas. Finally, we knockdown candidate TFs in primary and hiPSC-derived islets to validate their implication in regulating pancreatic GRNs.

## RESULTS

### Integrated Analysis of scRNA-seq Data Identifies 12 Human Adult Pancreatic Cell Types

Integrating multiple human adult pancreas scRNA-seq datasets can improve the power of scRNA-seq analyses to create a comprehensive human adult pancreas cell atlas. We set out to analyze and integrate five publicly available datasets covering a total of 35 non-diabetic, one type 1 diabetic and 15 type 2 diabetic individuals using Seurat v3.0 canonical correlation analysis (CCA) integration tools **(Figure 1A, Table S1)** (Segerstolpe et al. 2016; Y. J. Wang et al. 2016; Xin et al. 2016; Enge et al. 2017; Lawlor et al. 2017; Hafemeister and Satija 2019; Stuart et al. 2019). After filtering out low quality transcriptomes and data integration, uniform manifold approximation and projection for dimension reduction (UMAP) visualization revealed that 7393 cells localized into distinct clusters (**Figure 1B**). Cells from each original dataset localize together suggesting that the location of cells on the UMAP is not driven by the dataset of origin **(Figure 1B)**.

**Figure 1.**
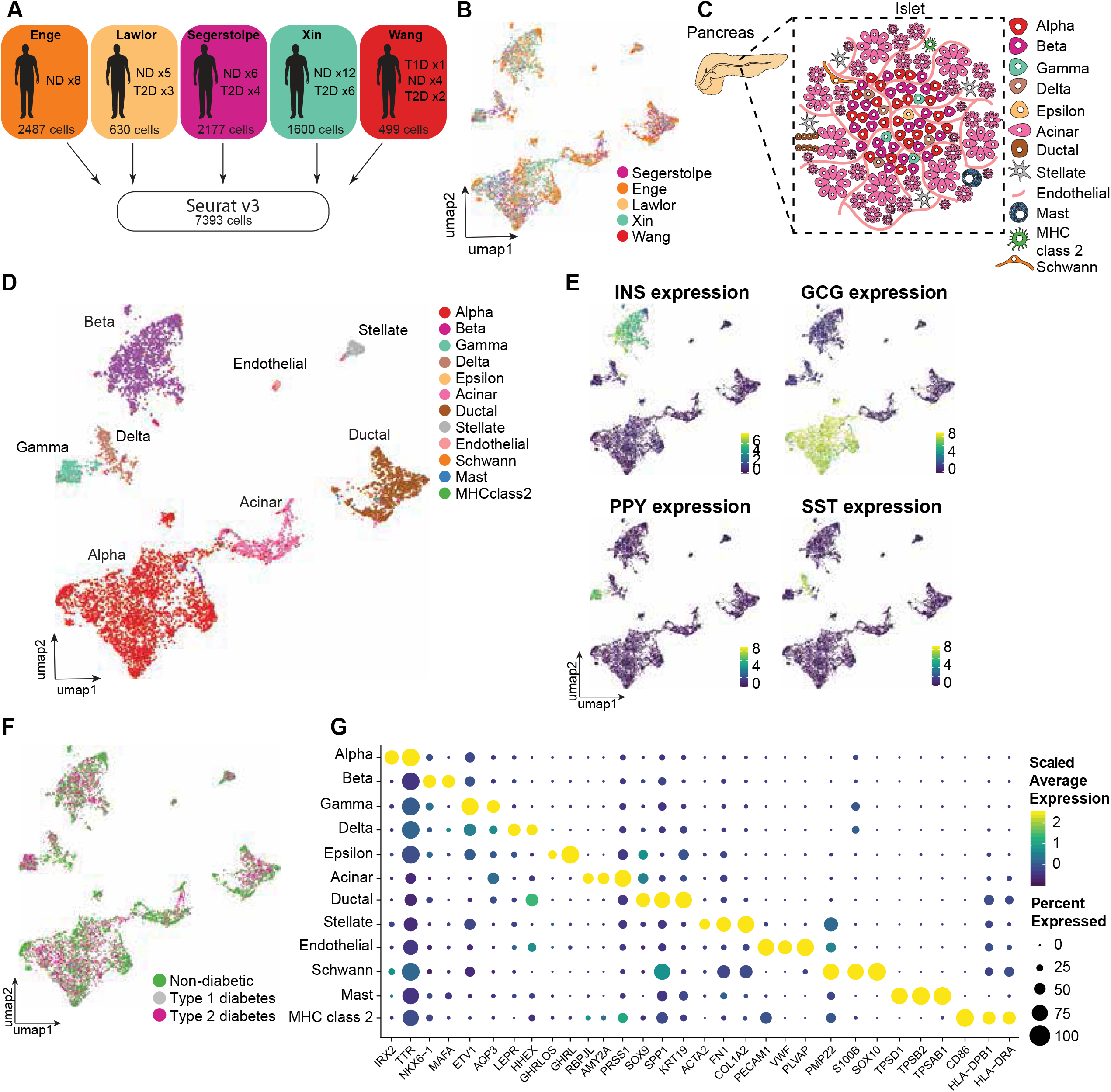
Integrated analysis of scRNA-seq identifies 12 pancreatic cell types. **(A)** Schematic of workflow used in this paper. Cells from five public datasets were processed uniformly from raw SRA files and integrated, resulting in one dataset of 7393 cells of 51 individuals. **(B)** Gene expression based UMAP of 7393 single cells annotated by dataset of origin. **(C)** Schematic overview of diverse cell types in the human adult pancreas. Created with BioRender.com **(D)** Integrated gene expression based UMAP of 7393 single cells annotated by cell type. **(E)** Gene expression based UMAP of 7393 single cells colored by non-integrated INS, GCG, PPY and SST expression. **(F)** Integrated gene expression based UMAP of 7393 single cells annotated by disease status. **(G)** Bubble plot showing various known marker genes across all annotated cell types. The bubble size is proportional to the percentage of cells that express a specific marker gene with the colour scale representing the average non-integrated scaled gene expression within the specific cell population.

We next sought to identify pancreatic cell types **(Figure 1C)**. Clustering analyses based on the expression of well-established cell type specific markers led to the identification of eight cell types in the human adult pancreas: beta, alpha, gamma, delta, acinar, ductal, stellate and endothelial cells **(Figure 1D, Table S2)**. UMAP visualization allowed the segregation of endocrine, exocrine and other lineages (**Figure S1A**). Beta cells grouped together, away from other clusters and were marked by INS (**Figure 1E**). Other distinct clusters corresponded to alpha, gamma and delta cells based on global transcriptional similarity and GCG, PPY and SST, and other markers, respectively **(Figure 1E, Table S2)**. Using a similar approach, we detected other, previously described, major pancreatic cell types including acinar, ductal, endothelial and stellate cells (**Figure 1D**). All cell types were detected in both non-diabetic and type 2 diabetic pancreases **(Figure 1F)**. Four additional rare cell populations, that cannot be robustly identified through clustering analyses, were identified manually by assessing GHRL (epsilon cells), TPS1AB (mast cells), CD86 (Major histocompatibility complex (MHC) class 2 cells) and SOX10 (schwann cells) **(Figure 1G)** (Wierup et al. 2002; Segerstolpe et al. 2016). These rare cell types often cluster with other common cell types. Importantly, our annotation largely recapitulated previous annotations **(Figure S1B,C)**. In summary, we reconstructed an integrated single-cell atlas of the human adult pancreas, and annotated 12 pancreatic cell types.

### Reconstruction of Gene Regulatory Networks in the Human Adult Pancreas

Next, we set out to comprehensively reconstruct GRNs for all pancreatic cell types from single-cell transcriptomic data, applying single-cell regulatory network inference and clustering (pySCENIC) (Aibar et al., 220; Sande et al. 2020). PySCENIC links *cis*-regulatory sequence information together with single-cell transcriptomes in three sequential steps by 1) co-expression analysis, 2) target gene motif and ChIP-seq track enrichment analysis, and 3) regulon activity evaluation **(Figure 2A)**. Each regulon consists of a TF with its predicted target genes (co-expressed genes with an enriched TF motif), altogether forming a regulon. pySCENIC identified 142 regulons that characterize the GRNs of the human adult pancreas **(Figure 2B,C, Table S3)**. Multiple regulons identified here as active in the pancreas correspond to TF binding motifs enriched in accessible chromatin in the pancreas, assessed by ATAC-seq in FACS-purified pancreatic cells (Arda et al. 2018), supporting the validity of the approach **(Figure S2A)**.

**Figure 2.**
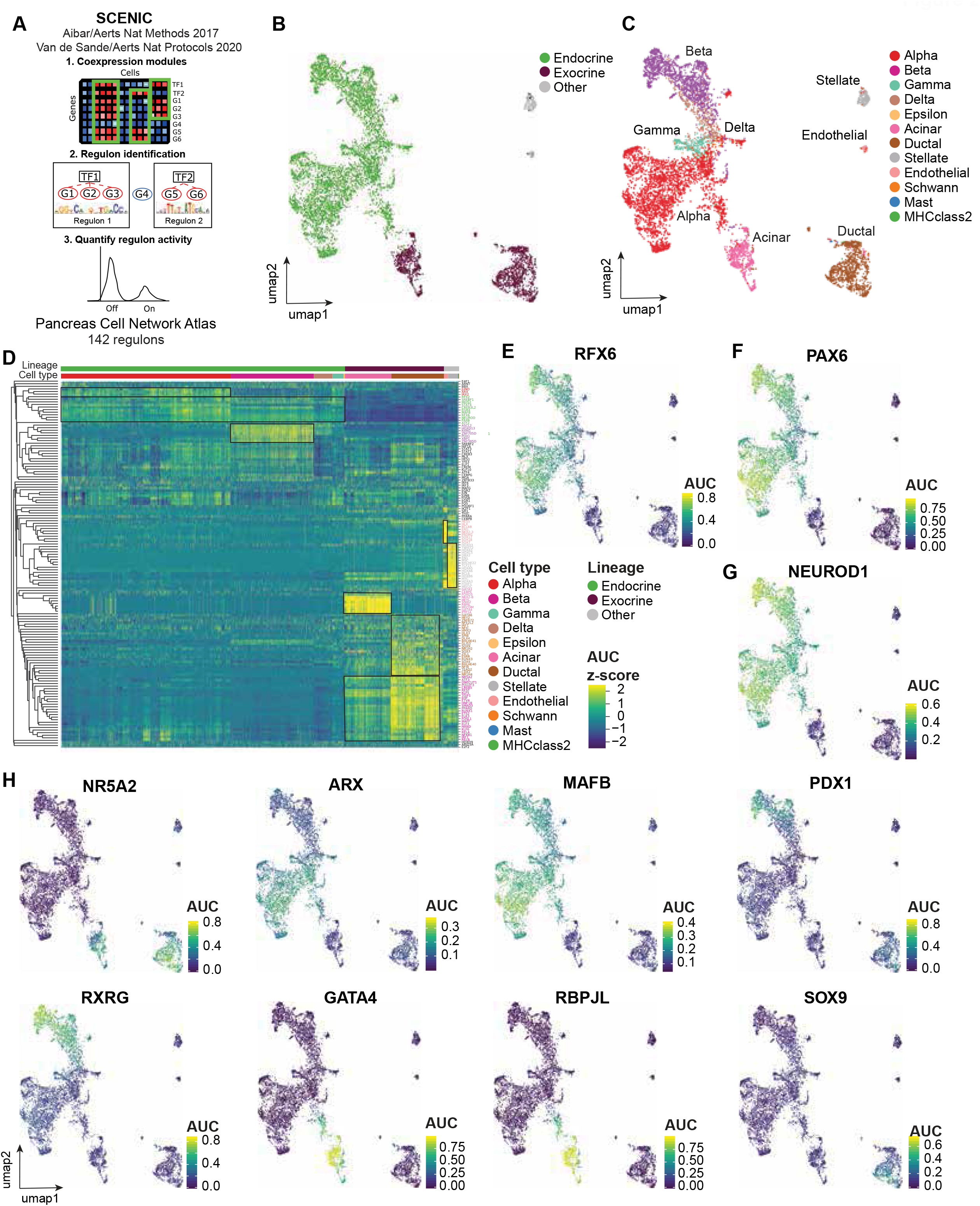
Reconstruction of gene regulatory networks in the human adult pancreas. **(A)** Schematic overview of the pySCENIC workflow used in this study. **(B-C)** Regulon activity based UMAP of 7393 single cells colored by pancreatic lineage **(B)** and cell type annotation **(C)**. Endocrine cells include alpha, beta, epsilon, gamma and delta cells. Exocrine cells include acinar and ductal cells. Other cells represent schwann, MHC class 2, mast, stellate and endothelial cells. **(D)** Heatmap based on unsupervised clustering of all 142 regulons (rows) for 4795 non-diabetic cells (columns). Color scaling is based on a z-score calculated based on the activity of each regulon. **(E-G)** Regulon activity based UMAP colored by the regulon activity of RFX6 **(E)**, PAX6 **(F)**, NEUROD1 **(G)**. **(H)** Regulon activity based UMAP colored by the regulon activity of NR5A2, ARX, MAFB, PDX1, RXRG, GATA4, RBPJL and SOX9.

UMAP visualization based on the activity of 142 regulons in non-diabetic and type 2 diabetic pancreata revealed groups of cells that differ from one another based on their regulatory activity **(Figure 2B,C)**. In particular, there are distinct regulatory states for exocrine and endocrine pancreatic lineages, stellate and endothelial cells (**Figure 2B,C**). Endocrine cell types clustered together, indicating shared regulatory states, while exocrine cell types formed two distinct clusters. Stellate and endothelial cells differed most from other cell types in their regulatory states. These results are consistent with previous analyses (Baron et al. 2016; Lawlor et al. 2017; Kumar and Vinod 2019) and are also in line with our findings based on gene expression analysis (**Figure 1D**). As expected, regulons active in endocrine cell types include RFX6, PAX6 and NEUROD1 **(Figure 2D-G)**. These TFs have reported roles in endocrine cell fate commitment and maintenance of cell identity throughout adult life (Smith et al. 2010; Mastracci et al. 2013; Piccand et al. 2014; Hart et al. 2013). Using iRegulon for visualization, many of the NEUROD1 target genes identified here have been previously linked to beta cell survival and function, such as SNAP25, TSPAN2, ELAVL4, PLCXD3 and NRNX1 (Mosedale et al. 2012; Hwang et al. 2016; Daraio et al. 2017; Juan-Mateu et al. 2017; Aljaibeji et al. 2019) **(Figure S2B)**. Interestingly, other NEUROD1 target genes are reported to be involved in cell fate specification during endocrine pancreas development such as PAX6, NKX2.2, INSM1 and HDAC6, suggesting an overlap in NEUROD1 target genes in adult life and embryonic development (Gierl et al. 2006; Hart et al. 2013; Haumaitre, Lenoir, and Scharfmann 2008). Clustering all cells based on the activity of all regulons identified regulatory modules (**Figure 2D**, black squares). In the exocrine pancreas, one regulatory module, containing NR5A2, was shared between acinar and ductal cells, although with a tendency for increased regulon activity in ductal cells **(Figure 2D,H)**. Other exocrine regulons included ONECUT1, REST and HNF1B, with reported roles in exocrine development (Nissim et al. 2016; Kropp, Zhu, and Gannon 2019) and the adult exocrine pancreas (Quilichini et al. 2019; Bray et al. 2020) **(Figure 2D)**. In summary, this analysis confirms the expected separation of exocrine and endocrine cells with distinct gene regulatory programs, and identifies known and novel candidate regulators of pancreas cell states.

Several regulatory modules are shared between different cell types within the endocrine and exocrine pancreas. Additionally, each cell type is defined by cell-type specific regulatory modules **(Figure 1D)**. In the endocrine pancreas, alpha and beta cells shared endocrine regulons (MAFB, MEIS2), whereas we observed distinct activities for ARX and IRX2 regulons in alpha cells and RXRG and PDX1 in beta cells **(Figure 2D,H)**, expanding previous findings (Kumar and Vinod 2019). Using iRegulon for visualization, PDX1 target genes include SLC6A17, PDIA6 and ABHD3, which have been reported to control insulin release (Eletto et al. 2016; Rorsman and Ashcroft 2018) **(Figure S2C)**. Interestingly, gamma and delta cells overlapped with alpha and beta cells, respectively, suggesting a shared regulatory state **(Figure 2C-D)**. This includes shared regulon activity for ARX in gamma and alpha and PDX1 in beta and delta cells **(Figure 2D,H)**, consistent with their reported expression in published scRNA-seq studies (Baron et al. 2016; Segerstolpe et al. 2016; Lawlor et al. 2017). GATA4 and RBPJL, known acinar-specific TFs (Carrasco et al. 2012; Masui et al. 2010), were highly active in acinar cells **(Figure 2H)**. Similarly, ductal cells were characterized by highly active SOX9 and POU2F3 regulons, in line with previous literature (Yamashita et al. 2017; Shroff et al. 2014) **(Figure 2D,H)**. In sum, this analysis confirms that alpha, beta, acinar and ductal cells are characterized by the activity of distinct combinations of active TFs that form gene regulatory modules.

In conclusion, the network approach recovers many of the expected regulators of pancreatic cellular identity allowing for the comprehensive characterization of the gene regulatory state of all major human adult pancreatic cell types.

### Prediction of Regulators of Endocrine Cell Identity in the Human Adult Pancreas

A comprehensive network analysis provides an opportunity to predict and identify critical regulators of cell identity. To identify regulons with highly cell type-specific activities within the human adult non-diabetic pancreas, we calculated regulon specificity scores (RSS) **(Table S4)** (Suo et al. 2018). The RSS utilizes Jensen*–*Shannon divergence to measure the similarity between the probability distribution of the regulon’s enrichment score and cell type annotation wherein outliers receive a higher RSS and are therefore considered cell type-specific (Suo et al. 2018). It can therefore be used to rank the activity of TFs within specific cell types.

Among the top regulons identified in alpha cells, we recover well known regulators of alpha and endocrine cell fate such as ARX, IRX2, PAX6, MAFB, NEUROD1 and RFX6 **(Figure 3A,B)** (Collombat et al. 2003; Artner et al. 2006; Delporte et al. 2008; Smith et al. 2010; Mastracci et al. 2013; Dorrell et al. 2011; Piccand et al. 2014). In addition, we identified JUND, EGR4, SREBF1 and STAT4 that have not yet been implicated in alpha cell identity. EGR1 (but not EGR4) has been shown to transcriptionally regulate GCG (Leung-Theung-Long et al. 2005) as well as the PDX1 promoter in beta cells (Eto, Kaur, and Thomas 2007). STAT4 and JUND have been described respectively in pancreatic tissue in general and in beta cells but not in alpha cells (Yu and Kim 2012; Good et al. 2019; K. Wang et al. 2021) **(Figure 3B,C)**. Interestingly, these TFs were also highly expressed in primary alpha cells found in the PANC-DB scRNA-seq dataset (Kaestner et al. 2019) **(Figure S3A)**. These TFs respond to the JNK and EGFR signaling pathways and may have important physiological functions. Both JUND and the JUND/JNK signaling pathway have been implicated in pancreatic cancer (Shin et al. 2009; Recio-Boiles et al. 2016). Immunocytochemistry of the human adult pancreas confirmed the presence of nuclear JUND in islets **(Figure 3Di)**. We also detected nuclear JUND protein in a subset of hiPSCs subjected to beta cell differentiation (**Figure 3E,F**). Surprisingly, we also detected JUND protein in ductal cells, despite lower JUND regulon activity in this cell type **(Figure 3C,Dii)**. Thus, protein expression does not necessarily mean that the TF is active in a given cell type. We also confirmed the expression of putative JUND target genes in primary alpha cells using the PANC-DB scRNA-seq dataset (Kaestner et al. 2019) **(Figure S3B)**. Nevertheless, these results show that JUND is present and active in a subset of pancreatic cell types in the human adult pancreas and hiPSC-derived islet cells. Altogether, this analysis predicts TFs active in human alpha cells, recovering known as well as new candidate TFs.

**Figure 3.**
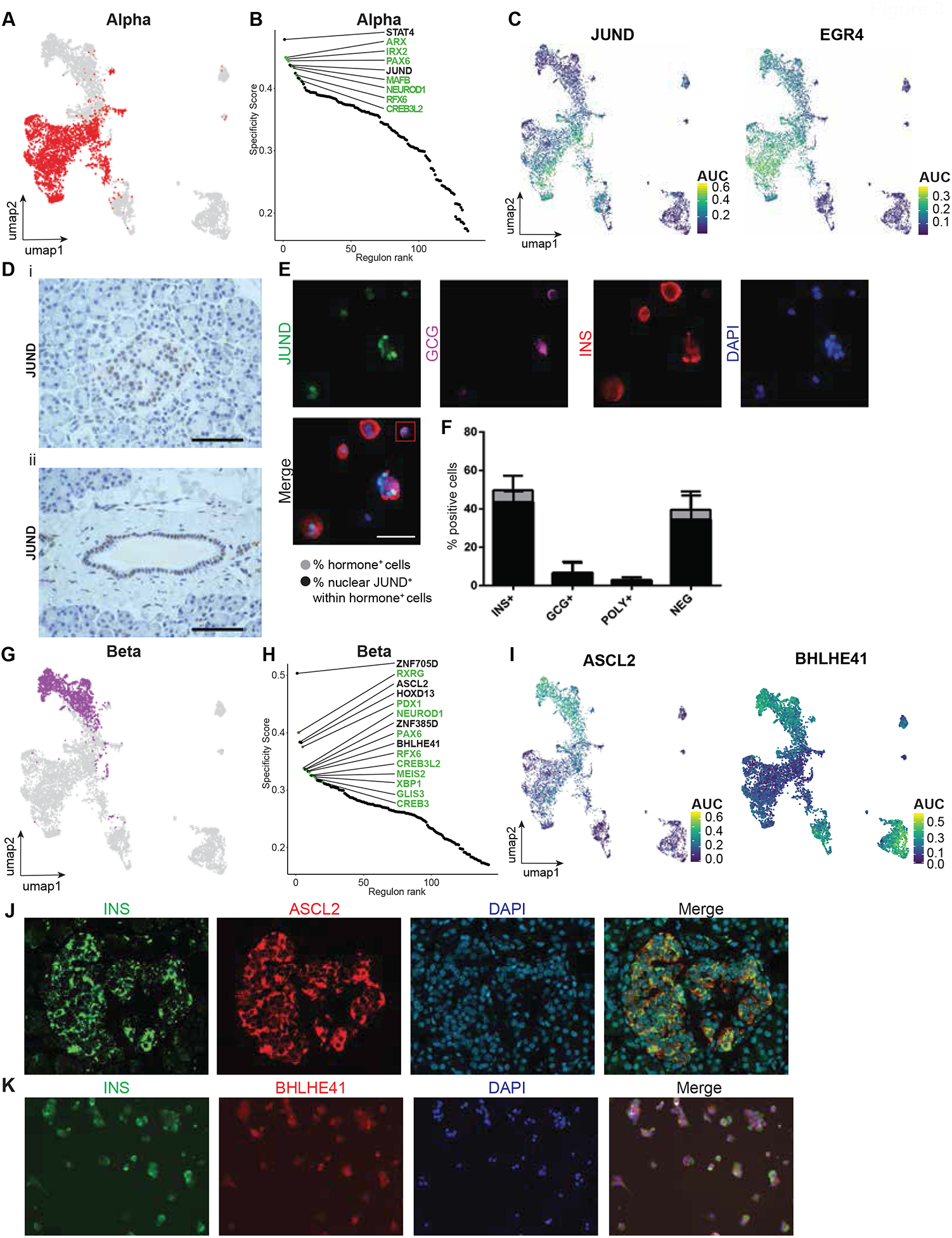
Prediction of regulators of endocrine cell identity in the human adult pancreas. **(A)** Regulon activity based UMAP of 7393 single cells with the alpha cell population highlighted. **(B)** Top 10 ranked regulons in non-diabetic alpha cells based on regulon specificity score (RSS). Known cell type specific regulons are colored green. **(C)** Regulon activity based UMAP colored by the regulon activity of JUND and EGR4 showing the cell type specificity of regulons. **(D)** Immunohistochemical evaluation of JUND in healthy human adult pancreata showing nuclear JUND expression in alpha, beta **(i)** and ductal **(ii)** cells. Representative images are shown (n = 3 individuals), Scale bar: 50 µm. **(E)** Immunofluorescence evaluation of JUND, INS, GCG and SST at stage 7 of hiPSC beta cell differentiation. Box indicates a GCG+ cell with nuclear JUND expression. Representative images examined for JUND (green), INS (red), GCG (purple) and DAPI (blue, nuclei counterstaining) are shown (n=3 beta cell differentiations). Scale bar: 50 µm. **(F)** The percentage of cells with nuclear JUND expression within the INS^+^, GCG^+^, polyhormonal and INS^-^/GCG^-^ cell populations. Results are shown as the normalized mean±s.d. (n=3 beta cell differentiations). **(G)** Regulon activity based UMAP of 7393 single cells with the beta cell population highlighted. **(H)** Top 10 ranked regulons in non-diabetic beta cells based on regulon specificity score (RSS). Known cell type specific regulons are colored green. **(I)** Regulon activity based UMAP colored by the regulon activity of ASCL2 and BHLHE41 showing the cell type specificity of regulons. **(J)** Immunohistochemical evaluation of ASCL2 in healthy human adult pancreata showing cytoplasmic ASCL2 expression in mainly beta cells. Representative images examined for INS (green), ASCL2 (red) and DAPI (blue, nuclei counterstaining) are shown (n = 3 individuals), Scale bar: 50 µm. **(K)** Immunohistochemical evaluation of BHLHE41 at stage 7 of hiPSC beta cell differentiation showing nuclear BHLHE41 expression in beta cells. Representative images examined for INS (green), BHLHE41 (red) and DAPI (blue, nuclei counterstaining) are shown (n = 2 beta cell differentiations), Scale bar: 50 µm.

Among the top regulons identified in beta cells, we retrieved well-known as well as new candidate regulators of beta and endocrine cell identity. Known TFs include RXRG, PDX1, NEUROD1, PAX6 and RFX6 (Zhou et al. 2008; Miyazaki et al. 2010; Smith et al. 2010; Mastracci and Sussel 2012; Hart et al. 2013; Piccand et al. 2014) (**Figure 3G-H)**. In addition, we found that ZNF705D, ASCL2, BHLHE41 and HOXD13 were highly ranked regulons (**Figure 3H-I)**. HOXD13 and BHLHE41 have been shown to be present in the exocrine pancreas (Cantile et al. 2009; Sato et al. 2012). We confirmed that these TFs were also expressed in primary beta cells of the PANC-DB scRNA-seq dataset (Kaestner et al. 2019) **(Figure S3C)**. Interestingly, ASCL2 has been reported to interact with β-catenin of the Wnt pathway; the latter has an established role in endocrine fate specification during *in vitro* differentiation (Schuijers et al. 2015; Vethe et al. 2019; Sharon et al. 2019). Many putative target genes of ASCL2 including PDX1, INS, ABCC8, FOXA1, KCNK16, FXYD2 are directly related to glucose sensing and beta cell identity, in line with the beta cell-specific regulatory activity of ASCL2 **(Figure S3D)** (Arystarkhova et al. 2013; Vierra et al. 2015; Hwang et al. 2016). FXYD2γa, a regulatory subunit of the Na+-K+-ATPase, is a transcript exclusively expressed in human beta cells (Flamez et al. 2010). Immunohistochemistry of human adult pancreas sections showed that ASCL2 is expressed in INS+ beta and islet cells **(Figure 3J)**. Surprisingly, ASCL2 was mainly localized to the cytoplasm **(Figure 3J)**, which is unexpected for TFs which tend to localize to the nucleus (Baranek, Sock and Wegner, 2005). Cytoplasmic localization of ASCL2 has been reported in the context of colon and breast cancer (R. Zhu et al. 2012; H. Xu et al. 2017). Immunohistochemistry of hiPSC-derived islet cells showed that BHLHE41 is expressed both in nuclei and cytoplasm of INS+ beta cells **(Figure 3K)**. We also confirmed the expression of putative BHLHE41 target genes using the PANC-DB scRNA-seq dataset (Kaestner et al. 2019) **(Figure S3E)**. These results implicate additional TFs including ASCL2 and BHLHE41 in the regulation of beta cell identity. They also illustrate the value of network analyses to increase our understanding of the biology of the human pancreas.

In summary, GRN analysis and regulon ranking allowed us to pinpoint both known and previously unknown candidate regulators of pancreatic endocrine cell identity, providing a resource for further investigation of their roles in cellular identity and function.

### BHLHE41 and JUND Depletion in hiPSC-Derived Islet Cells

To examine the effect of perturbation of candidate TFs in the human adult pancreas, we selected two TFs for targeting with siRNAs in hiPSC-derived islet cells, an *in vitro* pancreatic islet model (Balboa et al. 2022). JUND was identified as a regulator of oxidative stress and lipotoxicity in beta cells (K. Wang et al. 2021; Good et al. 2019) but its potential role in alpha cells remains unknown. BHLHE41 was of interest due to its role in circadian rhythm regulation (Hamaguchi et al. 2004) but its putative impact on cell identity in the human pancreas has not yet been investigated.

To gain insight into how candidate TF knockdown affects the transcriptome and cell identity, we performed scRNA-seq 72h post transfection **(Figure 4A)**. We confirmed that both JUND and BHLHE41 transcript levels were decreased in siRNA-transfected cells **(Figure 4B-C, S4A-B)**. BHLHE41 and JUND knockdown did not result in global gene expression changes suggesting that perturbation of these TFs is compatible with the maintenance of GRNs **(Figure 4D)**.

**Figure 4.**
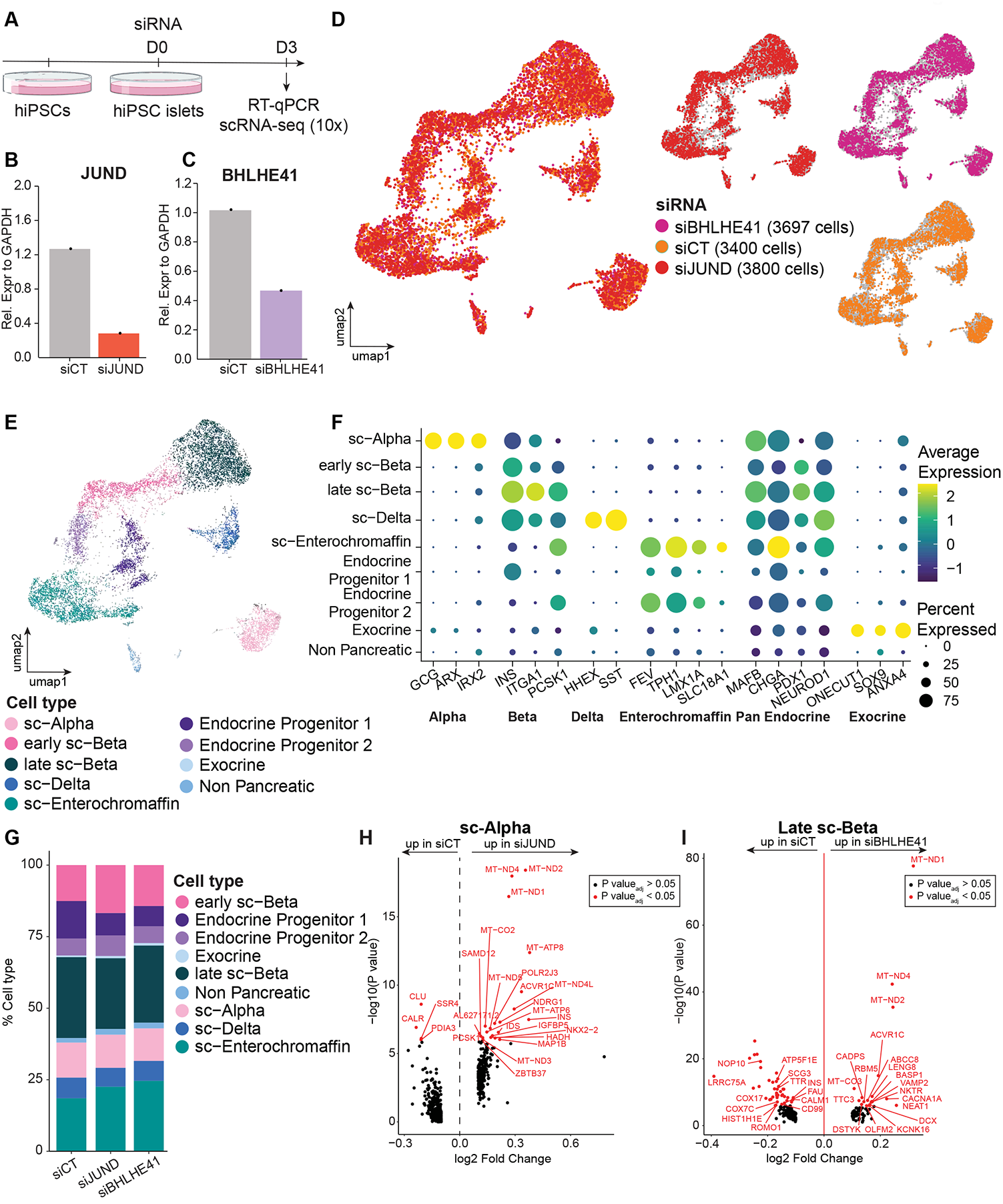
BHLHE41 and JUND depletion in hiPSC derived islet cells. **(A)** Scheme of siRNA mediated BHLHE41 and JUND knockdown in hiPSC islet cells. Created with BioRender.com **(B-C)** JUND **(B)** and BHLHE41 **(C)** transcript level 72h after siRNA transfection by RT-qPCR. Results are relative to the expression of GAPDH (arbitrary units) (n=1 beta cell differentiation). **(D-E)** Gene expression based UMAP of 10897 single cells annotated by siRNA transfection (D) and cell type (E). In D and E, boxes correspond to the 25th and 75th quartiles, horizontal lines to the median, and whiskers extend to 1.5 times the interquartile range. ***P<0.001 (Wilcoxon rank sum test with FDR correction compared to siCT). **(F)** Bubble plot showing the expression of various known marker genes across all annotated cell types. The bubble size is proportional to the percentage of cells that express a specific marker gene with the color scale representing the average scaled gene expression within the specific cell population. **(G)** Percentage of different cell types in each siRNA condition. **(H)** Volcano plot of differentially expressed genes in siCT and siJUND treated sc-alpha cells. Genes with an adjusted p-value < 0.05 are shown in red. Black represents genes that were not found to differ significantly between siRNA transfected groups. **(I)** Volcano plot of differentially expressed genes in siCT and siBHLHE41 transfected late sc-beta cells. Genes with an adjusted p-value < 0.05 are shown in red. Black represents genes that were not found to differ significantly between siRNA treated groups.

We next sought to annotate cell types within our sc-RNAseq dataset using clustering analyses and the expression of well-established cell type specific markers **(Figure 4E-F)**. This led to the identification of nine populations among the hiPSC islet cells: alpha, beta, delta, enterochromaffin, endocrine progenitor, exocrine and non-pancreatic cells **(Figure 4E)** in line with previous publications (Gonçalves et al. 2021; Rezania et al. 2014; Hrvatin et al. 2014; Z. Zhu et al. 2016; Augsornworawat et al. 2020; Hogrebe et al. 2021; Peterson et al. 2020; Veres et al. 2019; Balboa et al. 2022; H. Zhu et al. 2022).

To cross-reference our data with primary islets and published beta cell differentiation experiments, we integrated the dataset with previously described data (Balboa et al. 2022) **(Figure S4C-D)**. Most of the cell types identified in our dataset grouped together with previous datasets (Balboa et al. 2022; Krentz et al. 2018) and away from the primary human islet dataset (Xin et al. 2016) suggesting that our cell populations are transcriptionally similar to previously published datasets.

Next, we investigated whether BHLHE41 or JUND perturbation altered cell type proportions **(Figure 4G, Table S6)**. The number of sc-beta and sc-alpha cells remained unchanged upon either BHLHE41 and JUND perturbation, suggesting that TF knockdown did not result in specific loss of these cells **(Figure 4G)**. Sc-enterochromaffin cells are a known off-target population generated during the differentiation of hiPSCs towards the beta cell fate (Veres et al. 2019; Balboa et al. 2022). Recently, sc-enterochromaffin cells were found to more closely resemble a pre-beta cell population in the fetal pancreas (H. Zhu et al. 2022). Interestingly, both BHLHE41 and JUND deficiency tended to increase the number of sc-enterochromaffin cells (25% siBH, 23% siJU vs 19% siCT) **(Figure 4G)**.

Next, we examined which genes were affected by JUND or BHLHE41 depletion in hiPSC-derived islet cells by determining differentially expressed genes compared to the control condition (adjusted p-value <0.05, log2 fold change = 0.1). In sc-alpha cells, the expression of key beta cell genes INS, PCSK1, HADH and NKX2.2 (Flanagan et al. 2014) was increased upon JUND depletion **(Figure 4H)**. Additionally, GO analysis identified “insulin secretion” and “alpha cell to beta cell conversion” as top enriched pathways upon JUND depletion **(Figure S4E)**. Upon BHLHE41 depletion, sc-beta cells increased expression of multiple ion channels and proteins involved in endocrine cell electrical activity and granule exocytosis such ABCC8 (Meissner and Atwater 1976), CACNA1A (Braun et al. 2008), KCNK16 (Vierra et al. 2015) and VAMP2 (Regazzi et al. 1995). In contrast, the expression of key beta cell markers INS, SCG3 and TTR (Refai et al. 2005) decreased **(Figure 4I)**. GO analysis identified “Type 2 diabetes” as an enriched pathway upon BHLHE41 depletion **(Figure S4F)**. pySCENIC can be used to predict putative target genes of TFs **(Table S5)** (Aibar et al., 2017; Sande et al. 2020). We assessed if JUND and BHLHE41 knockdown altered the expression of their predicted target genes **(Table S5)**. The expression of the majority of BHLHE41 (10/12) and JUND (24/32) target genes in sc-alpha and sc-beta cells seemed to be unaffected upon siRNA treatment suggesting that the effect of TF knockdown was specific to a few genes **(Figure S4G-H)**.

### BHLHE41 Deficiency Induces Apoptosis in Human Adult Pancreatic Islets

GRNs were computed using primary pancreas scRNA-seq data (Xin et al. 2016; Y. J. Wang et al. 2016; Segerstolpe et al. 2016; Enge et al. 2017; Lawlor et al. 2017), but the validation experiments were carried out in hiPSC-derived islet cells. Since differences exist between hiPSC-derived islets and primary islets (Fantuzzi et al. 2022; Balboa et al. 2022), we next assessed whether JUND and BHLHE41 modulate cell identity in primary islets, using a knockdown approach. 72h after siRNA transfection into primary islet cells, we confirmed that JUND and BHLHE41 transcript levels were decreased **(Figure 5A-C, S5A-B)**. While JUND depletion did not affect human islet cell viability (Pearson correlation, R = -0.34, p = 0.23), reduced BHLHE41 transcript levels did correlate with increased apoptosis (Spearman correlation, R = -0.6, p = 0.025) **(Figure 5D-E, S5C)**. Interestingly, these results suggest that BHLHE41 promotes the survival of primary islets.

**Figure 5.**
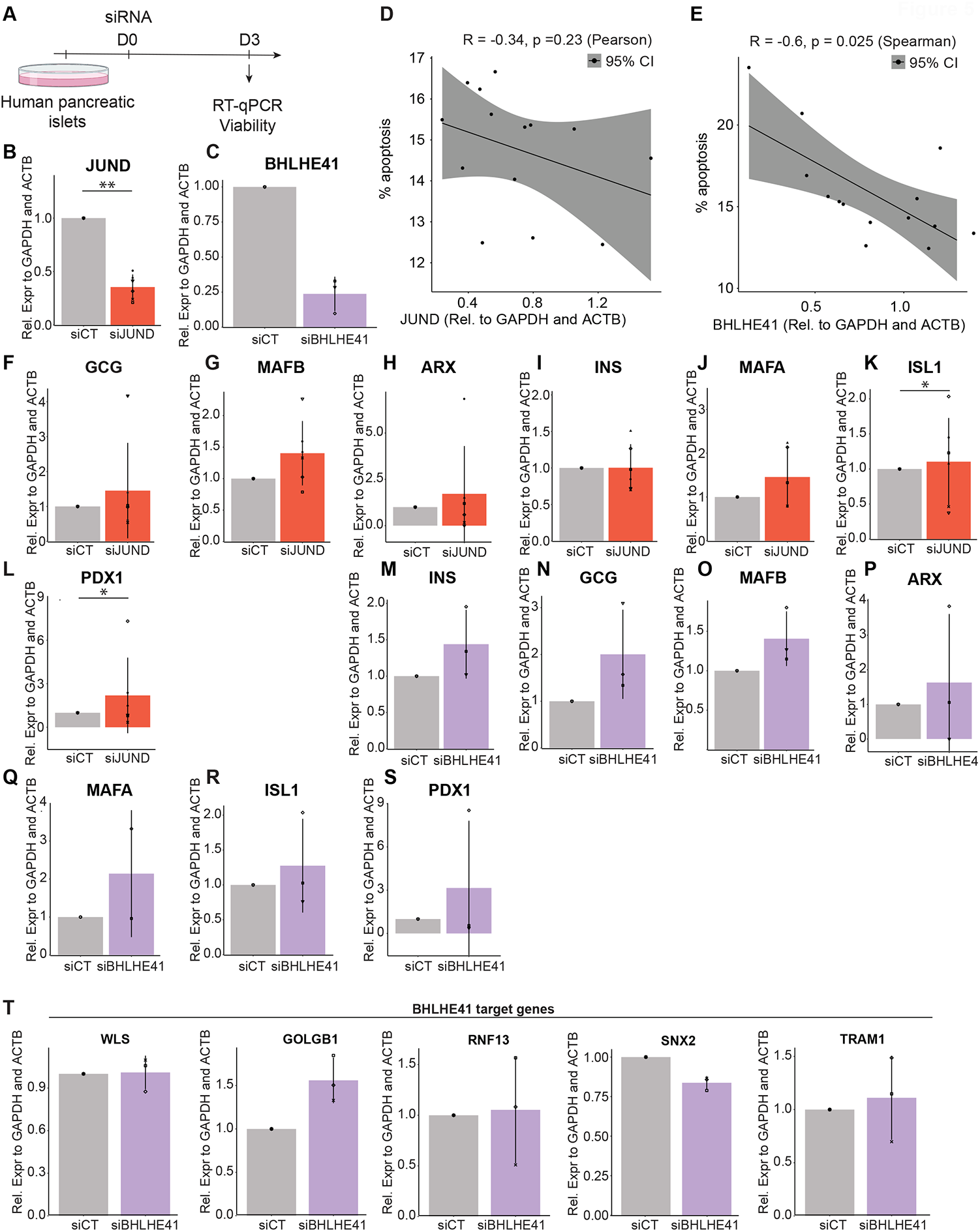
BHLHE41 deficiency induces apoptosis in human adult pancreatic islets. **(A)** Scheme of siRNA mediated BHLHE41 and JUND knockdown in primary pancreatic islets. Created with BioRender.com **(B-C)** JUND **(B)** and BHLHE41 **(C)** transcript levels 72 hours after siRNA transfection. **(D)** Pearson correlation between the percentage of apoptotic cells and JUND transcript level. **(E)** Spearman correlation between the percentage of apoptotic cells and BHLHE41 transcript level. **(F-L)** GCG **(F)**, MAFB **(G)**, ARX **(H)**, INS **(I)**, MAFA **(J)**, ISL1 **(K)** and PDX1 **(L)** transcript levels 72 hours after JUND siRNA transfection. n = 6 islet preparations **(M-S)** INS **(M)**, GCG **(N)**, MAFB **(O)**, ARX **(P)**, MAFA **(Q)**, ISL1 **(R)** and PDX1 **(S)** transcript levels 72 hours after BHLHE41 siRNA transfection. n = 3 islet preparations **(T)** Transcript level of BHLHE41 putative target genes WLS, GOLGB1, RNF13, SNX2 and TRAM1 72 hours post transfection in primary human islets. n = 3 islet preparations. *P<0.05, **P<0.01 (Wilcoxon rank sum test with FDR correction compared to siCT). Results are shown as the normalized mean±s.d. Each symbol represents one independent experiment.

JUND depletion had no significant effect on transcript levels of alpha cell marker genes GCG, MAFB and ARX, nor on beta cell marker genes INS and MAFA, while ISL1 and PDX1 transcript levels increased **(Figure 5F-L)**. Interestingly, BHLHE41 siRNA treatment tended to increase INS, GCG and MAFB transcript levels **(Figure 5M-O)**. We did not observe changes in ARX, MAFA, ISL1 and PDX1 transcript levels **(Figure 5P-S)**. We further assessed whether the expression of predicted BHLHE41 targets changed upon BHLHE41 knockdown. There was a trend for increased GOLGB1 expression, a component of the Golgi complex that is pivotal for proper insulin secretion **(Figure 5T)** (Stefan et al. 1987). Altogether, these data suggest that BHLHE41 deficiency induced apoptosis, increased INS, GCG and MAFB levels and altered transcript levels of a predicted target gene in primary islets.

### Prediction of Regulators of Exocrine Cell Identity in the Human Adult Pancreas

The comprehensive network analysis above also provides an opportunity to predict and identify regulators of exocrine cell identity.

We identified known and new TFs in acinar cells. Among the top acinar-specific regulons, we recovered well known regulators of acinar and exocrine cell identity such as PTF1A, RBPJL, GATA4 and NR5A2 (Ketola et al. 2004; Masui et al. 2010; Nissim et al. 2016; Sakikubo et al. 2018) **(Figure 6A/B)**. These findings are in line with a recent study that used single-nucleus RNA-seq on pancreatic acinar tissue (Tosti et al. 2021) and the expression of TFs in acinar cells of the PANC-DB scRNA-seq dataset (Kaestner et al. 2019) **(Figure S6A)**. Furthermore, we identified MECOM, HEYL and TGIF1 as highly ranked regulons **(Figure 6B/C)**. Interestingly, MECOM expression has been linked to acinar cell dedifferentiation which increases susceptibility to malignancy (Backx et al. 2021). The loss of TGIF1 has been linked to pancreatic ductal adenocarcinoma progression making further exploration of these regulons interesting in the context of cancer biology (Weng et al. 2019). Ectopic expression of *Tgif2* (but not *Tgif1*) reprograms mouse liver cells towards a pancreas progenitor state (Cerdá-Esteban et al. 2017). HEYL is a reported Notch signaling target gene in NGN3^+^ exocrine cells (Gomez et al. 2015). We confirmed nuclear expression of HEYL in human acinar and islet cells **(Figure 6Di/ii**, donor information can be found in **Table S1)** by immunohistochemistry, in agreement with elevated HEYL regulon activity in acinar cells **(Figure 6C)**.

**Figure 6.**
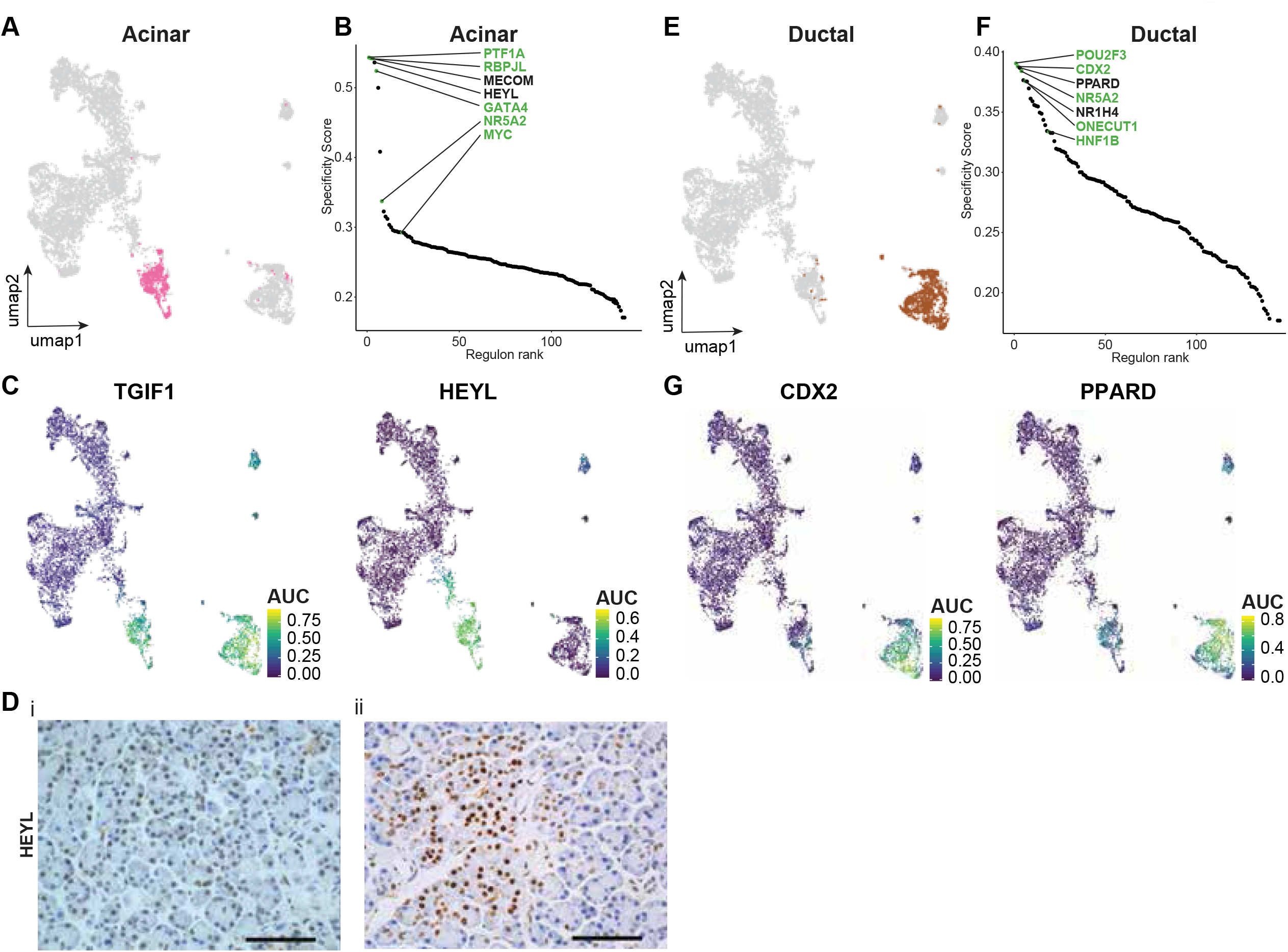
Prediction of regulators of exocrine cell identity in the human adult pancreas. **(A)** Regulon activity based UMAP of 7393 single cells with the acinar cell population highlighted. **(B)** Top 10 ranked regulons in non-diabetic acinar cells based on regulon specificity score (RSS). Known cell type specific regulons are colored green. **(C)** Regulon activity based UMAP colored by the regulon activity of TGIF1 and HEYL showing the cell type specificity of regulons. **(D)** Immunohistochemical evaluation of HEYL in healthy human adult pancreata showing nuclear HEYL expression in acinar **(i)** and islet **(ii)** cells. Representative images are shown (n = 3 individuals), Scale bar: 50 µm. **(E)** Regulon activity based UMAP of 7393 single cells with the ductal cell population highlighted. **(F)** Top 10 ranked regulons in non-diabetic beta cells based on regulon specificity score (RSS). Known cell type specific regulons are colored green. **(G)** Regulon activity based UMAP colored by the regulon activity of CDX2 and PPARD showing the cell type specificity of regulons.

Top ranked regulons in ductal cells included well known regulators of ductal and exocrine cell identity such as POU2F3, NR5A2 and HNF1B (Nissim et al. 2016; Yamashita et al. 2017; Quilichini et al. 2019) **(Figure 6E/F)**. In addition, we observed CDX2 and PPARD as highly specific ductal regulons **(Figure 6F/G)**. Both PPARD and CDX2 have been reported to be involved in human pancreatic ductal carcinoma, warranting further functional studies (K. Matsumoto et al. 2004; Coleman et al. 2013). We confirmed the expression of predicted TFs within ductal cells of the PANC-DB scRNA-seq dataset (Kaestner et al. 2019) **(Figure S6B)**.

In summary, GRN analysis and regulon ranking allowed us to pinpoint both known and predicted candidate regulators of pancreatic exocrine GRNs and cell identity. Specifically, we identified HEYL as a candidate TF that might be implicated in regulating acinar cell identity, warranting further investigation.

### An Interactive Online Resource for Visualization of the Human Adult Pancreas Cell Network Atlas

To enable users to easily navigate the human pancreatic cell network atlas, we provide a loom file that allows for the visualization and exploration of the data using the web-based portal SCope (Davie et al. 2018) **(.loom file and tutorial available at** http://scope.aertslab.org/#/PancreasAtlas/*/welcome **and** https://github.com/pasquelab/scPancreasAtlas). Features such as cell type annotation as defined in this paper, gene expression and regulon activity, can be explored on the regulon and gene expression based UMAPs. This resource enables users to select and visualize up to three genes or regulons simultaneously and select subsets of cells for downstream analyses. Target genes of a specific regulon can be downloaded to facilitate further exploration, for example in iRegulon or gene ontology analysis (Janky et al. 2014). A list of predicted target genes of all 142 regulons can also be found in **Table S5**. Furthermore, a list of target genes can be manually defined to compute the activity of a custom regulon. This resource can be used to further study cell identity, GRNs and gene regulation in the context of the pancreas.

## DISCUSSION

In this resource, we take advantage of integration strategies and new computational tools to reconstruct an integrated cell and GRN atlas of the human adult pancreas from single-cell transcriptome data. This approach provides a comprehensive analysis of the gene regulatory logic underlying cellular identity in the human adult pancreas in a broad range of individuals, limiting the influence of inter-donor variability. We recovered known regulators of pancreatic cell identity and uncovered predicted candidate regulators of cell identity that can be further investigated for their roles in cellular identity and function. By validating regulon analyses and creating an easily accessible interactive online resource which allows for the exploration of the gene regulatory state of 7393 cells from 51 individuals, this approach extends beyond previous gene regulatory studies in the human adult pancreas (Kumar and Vinod 2019).

The present analysis identified regulators of pancreatic development, function and survival that are known to be critical in humans because loss-of-gene function causes pancreatic agenesis or young onset diabetes. For example, PTF1A and GATA4, whose loss of function are linked to pancreatic agenesis and neonatal diabetes (Sellick et al. 2004; Weedon et al. 2014; Shaw-Smith et al. 2014), were among the top acinar-specific regulons **(Figure 6)**. In addition, monogenic diabetes related genes PDX1 (Nicolino et al. 2010), NEUROD1 (Rubio-Cabezas et al. 2010), PAX6 (Solomon et al. 2009), RFX6 (Smith et al. 2010; Patel et al. 2017) and GLIS3 (Senée et al. 2006) were among the top beta cell-specific regulons **(Figure 3 and Table S4**).

Several top regulons in endocrine but not exocrine cells are TFs involved in endoplasmic reticulum stress signaling (**Table S4**). CREB3 and CREB3L2 are non-canonical endoplasmic reticulum stress transducers that are induced in human islets and clonal beta cells upon exposure to the saturated fatty acid palmitate (Cnop et al. 2014). Interestingly, SREBF1 and -2 undergo similar endoplasmic reticulum exit and proteolytic processing in the Golgi as these endoplasmic reticulum stress transducers, but they do so in response to changes in endoplasmic reticulum cholesterol content; both also have high regulon activity in alpha and beta cells. XBP1 is abundantly expressed in the exocrine and endocrine pancreas (Cnop et al. 2017), but the XBP1 regulon has its highest specificity in beta cells. ATF3 and ATF4 are TFs that are activated upon eIF2α phosphorylation, an endoplasmic reticulum stress response pathway to which no less than 5 monogenic forms of diabetes belong (Eizirik, Pasquali, and Cnop 2020). Our data underscore the importance of these TFs for endocrine pancreatic cell identity.

Disruptions of circadian clock genes within the pancreas have been linked to impaired glucose tolerance and type 2 diabetes (Boden, Chen, and Polansky 1999; Pappa et al. 2013; Alvarez-Dominguez et al. 2020). The mammalian circadian clock consists of transcriptional oscillators that coordinate behavior and metabolism within a 24h light-dark cycle. The core loop consists of BMAL1-CLOCK that activate clock-controlled genes, including PER and CRY genes, which in turn form complexes that inhibit CLOCK-BMAL1 mediated transcription upon entry in the nucleus (Langmesser et al. 2008). Here, we investigated the role of BHLHE41, a transcriptional repressor of CLOCK-BMAL1 that in turn activates clock target genes (Fujimoto et al. 2007; Y. He et al. 2009). We found that BHLHE41 deficiency induced apoptosis in human islets. This finding is consistent with reports of apoptosis caused by disrupting the circadian rhythm of diabetic rats (Gale et al. 2011). The high level of redundancy between different clock genes could help explain the limited transcriptional effects of the depletion of a single clock gene (Wu et al. 2021). It is speculated that this genetic and functional redundancy of clock genes ensures tight control and entrainment of circadian rhythm (Oster et al. 2002, 2003; Hogenesch and Herzog 2011).

Given that we predict regulators of cell identity in the human pancreas, it will be interesting to expand this analysis to embryonic development of the pancreas (Cao et al. 2020; Han et al. 2020; la O Sean et al. 2022; Gonçalves et al. 2021; Y. Xu et al. 2022). Our work may also be beneficial in guiding the *in vitro* differentiation of pancreatic cell types and minimize the emergence of off-target cell populations such as sc-enterochromaffin cells. For example, the emergence of SST-positive cells together with beta-like cells at the end of *in vitro* differentiation could be explained by the overlap in regulatory states between beta and delta cells (Baron et al. 2016). A better understanding of the regulatory logic underlying the control of beta cell fate through these GRN analyses may help improve or facilitate future applications in regenerative medicine (Pagliuca et al. 2014; Rezania et al. 2014; Nostro et al. 2015; Russ et al. 2015; Baeyens et al. 2018). Alternatively, many TFs such as ASCL2, MECOM, PPARD, GATA6 and CDX2 are linked to pancreatic cancer making the exploration of GRNs interesting in the context of cancer biology (K. Matsumoto et al. 2004; R. Zhu et al. 2012; Coleman et al. 2013; Hoang et al. 2016; H. Xu et al. 2017; Weng et al. 2019; Brunton et al. 2020). Recent reports have stratified type 2 diabetes patients based on age at diagnosis, BMI, HbA1c and insulin secretion and sensitivity, and identified subtypes with different genetic predisposition, treatment response, disease progression and complication rates (Ahlqvist et al. 2018). Hence, it would be interesting to assess differences in gene regulatory state and gene expression profiles of alpha and beta cells between different type 2 diabetic subgroups.

Taken together, our GRN atlas, containing 51 individuals, provides a valuable resource for future studies on human pancreas homeostasis, donor variability, development, and disease including type 2 diabetes and pancreatic cancer. Finally, our results provide new insights into the activity of TFs and gene regulation in the human adult pancreas from a gene regulatory perspective.

### Limitations of the study

It is important to note that pySCENIC is a stochastic algorithm that does not produce precisely the same regulons for repeated runs, limiting reproducibility when comparing different datasets and when SCENIC is used multiple times on the same dataset (Huynh-Thu et al. 2010; Sande et al. 2020). To mitigate this uncertainty, we ran the full pySCENIC pipeline five times and only kept consistent regulons with the highest regulon activity. Alternatively, SCENIC could be automated to be run hundreds of times to further mitigate stochasticity (Sande et al. 2020). The performance of pySCENIC and other GRN inference methods suffers due to the large amount of drop-out events in scRNA-seq data warranting caution when interpreting results (Chen and Mar 2018). This could explain the absence of well-established pancreas TFs such as MAFA (Olbrot et al. 2002), MNX1 (Flanagan et al. 2014), NEUROG3 (Krentz et al. 2017), FOXA2 (Lee et al. 2019) and NKX2-2 (Mastracci et al. 2011) in this analysis. Nevertheless, in support of the validity of our findings, ATAC-seq, the literature and immunohistochemistry of human pancreas sections corroborate several pySCENIC predictions reported here such as BHLHE41, JUND and HEYL (Gomez et al. 2015; Good et al. 2019; K. Wang et al. 2021). Chen and colleagues underline the importance of using large sample sizes to derive the most accurate network inference possible (Chen and Mar 2018), highlighting the importance of dataset integration to increase the number of cells analyzed. In the future, it will be interesting to extend these analyses to include many more cells and patients. Despite current caveats, GRN analysis has enabled the capture of biological relevant information (Butte et al. 2000).

In this work, we tested the effect of BHLHE41 and JUND knockdown in both hiPSC-derived and primary islets. The absence of complete BHLHE41 and JUND depletion could explain the limited effect on candidate target genes. CRISPR knockout approaches could be used in the future to mitigate this limitation (Bevacqua et al. 2021). Additionally, we cannot exclude that the dispersion of cells and 2D culture prior to siRNA transfection affects islet behavior (Hopcroft, Mason, and Scott 1985).

One additional limitation of this study is the assumption that all TFs bind their binding motifs in the promoters of expressed genes. However, TF binding can be restricted to a subset of TF motifs in the genome due to influence of chromatin processes including the presence of nucleosomes as well as DNA methylation. Therefore, additional approaches such as single cell multi-omics that capture additional layers of genome regulation will be helpful to increase our understanding of gene regulation in the context of the human pancreas. Recently developed computational tools including SCENIC+ and CellOracle could help towards this goal (González-Blas et al. 2022; Kamimoto, Hoffmann, and Morris 2020).

## ACKNOWLEDGEMENTS

We thank Stein Aerts, Kristofer Davie and the Stein Aerts lab for discussions and creating the permanent SCope link, Shengbao Suo for sharing the MATLAB script for calculating the regulon specificity score, the Flemish Supercomputer Center (VSC) and Leuven Stem Cell Institute. We thank Stein Aerts for feedback on the manuscript. Research in the Pasque laboratory was supported by the Research Foundation– Flanders (FWO; Odysseus Return Grant G0F7716N to V.P.; FWO grants G0C9320N and G0B4420N to V.P.), the KU Leuven Research Fund (BOFZAP starting grant StG/15/021BF to V.P. and C1 grant C14/21/119 to V.P.), and FWO SB Ph.D. Fellowship to L.V. (1S29419N), and by the Pandarome project 40007487 (G0I7822N) (funded by the FWO and F.R.S.-FNRS) under the Excellence of Science (EOS) programme to M.C. and V.P. Research in the Cnop lab was also funded by the Fonds National de la Recherche Scientifique (FNRS) to M.C. and F.F., the Marie Skłodowska-Curie Actions Fellowship from the European Union’s Horizon 2020 research and innovation programme under the Marie Skłodowska-Curie grant agreement No 801505 to T.S., the Walloon Region SPW-EERWin2Wal project BetaSource and the Francophone Foundation for Diabetes Research (sponsored by the French Diabetes Federation, Abbott, Eli Lilly, Merck Sharp & Dohme and Novo Nordisk) to M.C., the European Foundation for the Study of Diabetes/Boehringer Ingelheim European Research Programme on ‘Multi-System Challenges in Diabetes’ to M.C. and F.F. and the Innovative Medicines Initiative 2 Joint Undertaking under grant agreement No 115797 (INNODIA; this Joint Undertaking receives support from the Union’s Horizon 2020 research and innovation program and “EFPIA” (European Federation of Pharmaceutical Industries Associations), “JDRF” (Juvenile Diabetes Research Foundation), and “The Leona M. and Harry B. Helmsley Charitable Trust”). This manuscript used data acquired from the Human Pancreas Analysis Program (HPAP-RRID:SCR_016202) Database (https://hpap.pmacs.upenn.edu), a Human Islet Research Network (RRID:SCR_014393) consortium (UC4-DK-112217, U01-DK-123594, UC4-DK-112232, and U01-DK-123716)

## AUTHOR CONTRIBUTIONS

Conceived project, VP, LV. Performed experiments, FF, AAS, MVH. Analyses, FF, LV, AAS, MVH, TH, AR, TH, TS, TR. scRNA-seq analyses, LV, XY, AJ, JC. ATAC-seq analysis, LV. Resources, MC, JKC. Wrote manuscript, LV, VP and MC with input from all authors. Supervision, VP.

## METHODS

### Key Resources Table

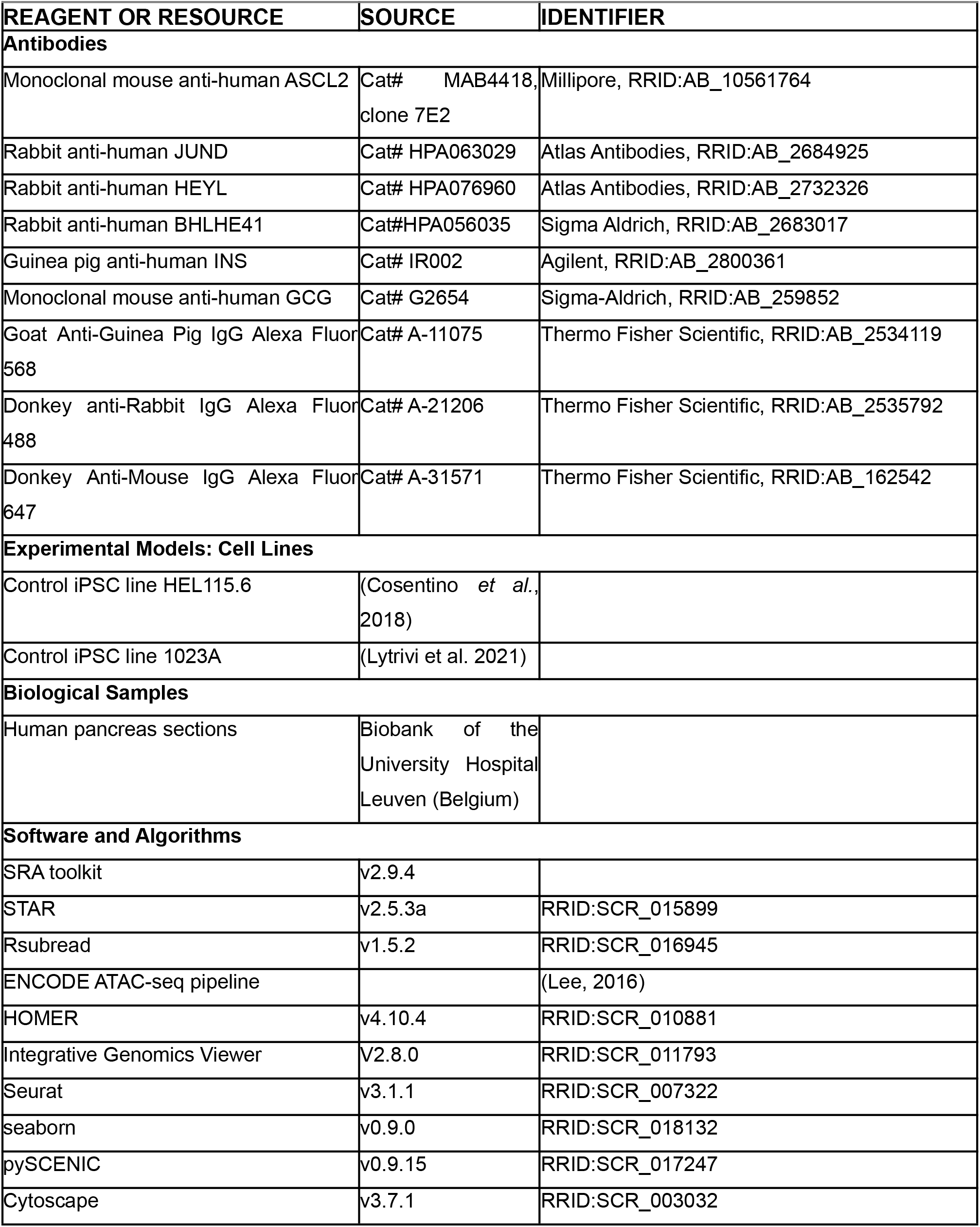
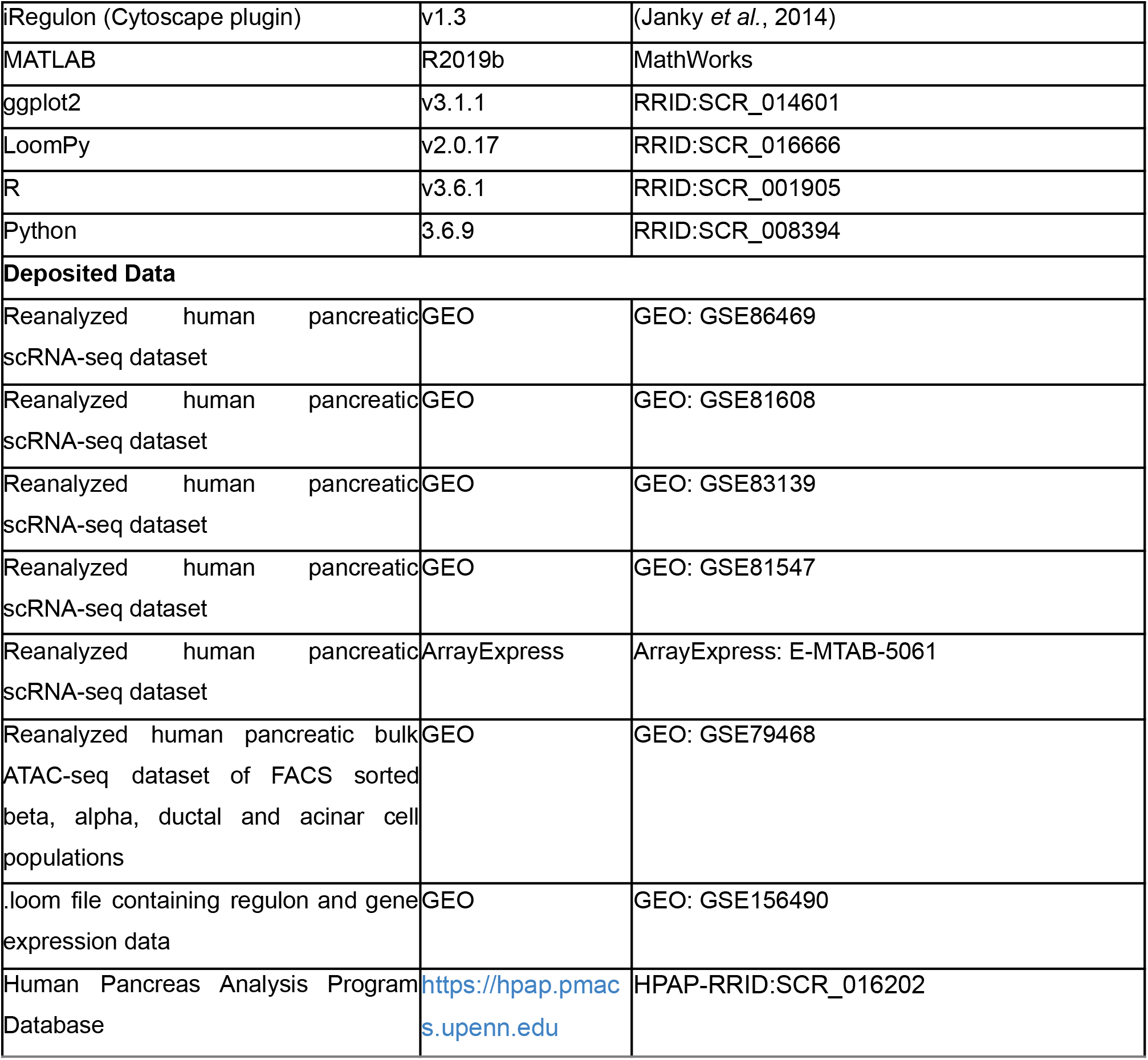

### Resource availability

#### Lead Contact

Further information and requests for resources should be directed to and will be fulfilled by the Lead Contact, Vincent Pasque.

#### Materials Availability

This study did not generate new unique reagents.

#### Data and Code Availability

Regulon data, raw and integrated gene expression matrices and the .loom file are available in the Gene Expression Omnibus (GEO) repository under accession code GSE156490 [https://www.ncbi.nlm.nih.gov/geo/query/acc.cgi?&acc=GSE156490]. A SCope tutorial is available at https://github.com/pasquelab/scPancreasAtlas. This study did not generate any new software; questions about data analysis should be directed to the Lead Contact, Vincent Pasque. The reviewer tokens for this GEO repository is cfsjciumfferjkd.

### Method Details

#### Motif discovery of bulk ATAC-seq data

Paired-end raw reads for bulk ATAC-seq (see Key Resource Table) were downloaded from SRA using SRA toolkit (v2.9.4). Reads were aligned and further analyzed using the ENCODE ATAC-seq pipeline with default parameters using the ENCODE human reference genome GRCh38.15 (Lee, 2016). Bed files containing the global open chromatin landscape of Adult Alpha (Alpha_1; EA4 and Alpha_2; EA28), Beta (Beta_1; EA5 and Beta_2; EA29), Acinar (Acinar_1; EA7 and Acinar_2; EA27) and ductal (EA11) cells or cell type specific differentially accessible regions were used as input for motif discovery by HOMER (v4.10.4) using the ‘findMotifsGenome.pl’ with options using hg38 with size given (Heinz *et al*., 2010). The TFs whose motifs identified by HOMER correspond with TFs identified by pySCENIC are visualized in Figure S2A.

#### Analysis of publicly available scRNA-seq data

Raw reads for five publicly available scRNA-seq datasets (see Key Resource Table) were downloaded from SRA using SRA toolkit (v2.9.4). Afterwards, reads were aligned to the human reference genome GRCh38.95 using STAR (v2.5.3a) with default parameters followed by the conversion to the coordinate sorted BAM format. Next, the featureCounts command from the “Rsubread” (v1.5.2) package in R (v3.6.1) was used to assign mapped reads to genomic features. Low quality transcriptomes with a mitochondrial contamination greater than 5% and less than 200 expressed genes per cell were excluded from subsequent analyses. The resulting raw count matrix was batch corrected using the FindIntegrationAnchors and IntegrateData functions from the “Seurat” package (v3.1.1) after which subsequent analyses were carried out in the R package “Seurat” (v3.1.4). Gene expression was used to cluster all 7393 cells with UMAP, using Seurat’s function RunUMAP.

Clusters for cell type annotation were defined using Seurat’s shared nearest neighbor algorithm FindClusters function after which differential expression analysis was performed using Wilcoxon’s rank sum test with a minimum cutoff of 0.25 average log fold change and min.pct of 0.25.

#### pySCENIC

GRNs were inferred using pySCENIC (python implementation of SCENIC, v0.9.15) in Python version 3.6.9 (Aibar et al., 2017). Integrated read counts were used as input to run GENIE3 (Huynh-Thu et al. 2010) which is part of arboreto (v0.1.5). GRNs were subsequently inferred using pySCENIC with the hg38_refseq-r80 motif database and default settings. To control for the stochasticity, which is inherent to pySCENIC, a consensus GRN was generated by merging results from five repeat pySCENIC runs. If regulons were identified in multiple pySCENIC runs, only the regulon with the highest AUC value was retained. Regulon activity represented by AUCell values was used to cluster all 7393 cells with UMAP, using Seurat’s RunUMAP function.

All 142 regulons within non-diabetic cell types were visualized using the ‘clustermap’ function of the Python package “seaborn” (v0.9.0). The z-score for each regulon across all cells was calculated using the z-score parameter of the seaborn ‘clustermap’ function.

Extended analysis of the target genes of specific regulons was conducted in Cytoscape (v3.7.1) using the iRegulon application (v1.3). The list of target genes of a specific regulon was downloaded from the loom file through the SCope platform (https://github.com/pasquelab/scPancreasAtlas) (Davie et al. 2018).

#### Regulon ranking

To quantify the cell-type specificity of a regulon, we utilized an entropy-based strategy as described previously (Suo *et al*., 2018) using the AUCell matrix as input in MATLAB R2019b. The top 10 most specific regulons were subsequently visualized using the R package ggplot2 (v3.1.1). The complete regulon ranking list is available in Table S2.

#### HPAP scRNA-seq data processing and analysis

Raw fastq files of single cell RNA-seq data were obtained from the portal of Human Pancreas Analysis Program (https://hpap.pmacs.upenn.edu). We analyzed 12 T2D donors (HPAP-058, 088, 065, 051, 081, 083, 085, 057, 091, 079, 070, 061) and 32 ND donors (042, 044, 039, 047, 034, 038, 043, 024, 072, 050, 019, 029, 036, 026, 082, 045, 052, 049, 027, 056, 035, 037, 040, 059, 075, 022, 054, 074, 063, 077, 093, 053). Cell Ranger (10x Genomics; v6.1.2) was used for alignment (reference genome hg38) and filtering of sequencing reads. For each sample, decontamination of the ambient mRNA was done with SoupX (v1.6.1) by using the automatic selection of the background genes to adjust the gene profiles. The new single-cell gene expression profiles were imported into Seurat (v4.1.1) for quality control. We kept genes expressed in at least three cells that express at least 200 genes. Potential doublet cells were evaluated by R package scDbIFinder (v3.16) and removed. Further filtering was performed by criteria of nFeature_RNA < 9000, percent of mitochondrial genes <10 and nCount_RNA <10000. We obtained 78,927 single cells for downstream analysis. We next used the seurat’s SCTransform (SCT) function to measure the differences in sequencing depth per cell and normalize the counts by removing the variation due to sequencing depth (nUMIs). Top 3000 of the variable features were selected to perform dimensionality reduction by PCA and UMAP embedding. Lastly, R package scSorter (v0.0.2) was used to assign cells to known cell type based on the marker genes. We extract the “alpha”, “beta”, “acinar” and “ductal” cells to check the top 10 regulons predicted by SCENIC analysis in the present study.

#### scRNA-seq sample preparation

hiPSC islets were washed with Versene (Thermo Fisher Scientific, 15040066) and incubated with Accutase (Sigma-Aldrich, A6964-100ML) for 10 min at 37°C. Dissociation was stopped by the addition of PBS supplemented with 1% Bovine Serum Albumin (BSA). Dissociated cells were washed twice in PBS (1%BSA) buffer, centrifuged at 300 rcf for 5 min, filtered using Flowmi 40µm tip strainer (Bel-Art, H13680-0040), counted and adjusted to 1 × 10^6^ cells ml–1 cells in PBS (1%BSA) for encapsulation. Cell count and viability were measured using the Luna-Fl automated Fluorescence Cell Counter (Logos Biosystems).

For each siRNA treatment, 6000 cells were loaded onto the 10X Chromium Single Cell Platform (10X Genomics) using the Next GEM Single Cell 3’ library and Gel Bead Kit (v3.1 chemistry) according to manufacturer’s instructions (10x User Guide CG000204, Revision D). RNA quality was assessed using Tapestation (Agilent). Libraries were quantified using Tapestation (Agilent), the Qubit 2.0 (ThermoFisher Scientific) and KAPA Library Quantification Kit for Illumina Platform (KAPA Biosystems) before pooling. Libraries were pooled in equimolar amounts for paired-end sequencing on an Illumina NextSeq 2000 instrument to yield ∼168 million (range 147–195 million) 100-bp-long reads on average per sample.

#### scRNA-seq analysis

Raw data was processed using the 10x Genomics CellRanger software (v4.0.0) and mapped to the pre-built human reference genome GRCh38-2020-A. Contamination by background RNA from disrupted cells was estimated and corrected for using SoupX(Young and Behjati 2020) with Seurat identified clusters, and known cell type specific marker genes (GCG, TTR; INS, IAPP; SST; PPY; GHRL; CPA1, CLPS, CPA2, REG1A, CELA3A, CTRB1, CTRB2, PRSS2; KRT19, VTCN1). Cells with less than 200 expressed genes and more than 25% mitochondrial reads were excluded from future analyses. DoubletFinder (v2.0.3) (McGinnis, Murrow, and Gartner 2019) was used with default settings to identify and remove potential doublets. The resulting counts were normalized, scaled and analyzed for principal component analysis (PCA) and uniform manifold approximation and projection (UMAP) on the first 20 principle components (PCs) using 2000 variable genes using Seurat (v3.1.1). Clusters for cell type annotation were defined using Seurat’s shared nearest neighbor algorithm as defined above.

To compare our dataset to the sc-derived and primary cell populations described in (Balboa et al. 2022), we downloaded the countmatrix and metadata files from the Single Cell Portal from the Broad Institute (SCP1526). The resulting Seurat object was integrated with our dataset using the FindIntegrationAnchors and IntegrateData functions from the “Seurat” package (v3.1.1) after which subsequent analyses were carried out in the R package “Seurat” (v3.1.4). Gene expression was used to cluster all 57158 cells with UMAP, using Seurat’s function RunUMAP.

#### Human islets

Human islets (seven non-diabetic donors, age 43 ± 15 years, BMI 30 ± 6 kg/m2 and one type 2 diabetic donor, age 66 years, BMI 36.5 kg/m2) were isolated by enzymatic digestion and density gradient purification in Lille or Edmonton, with the approval of the local Ethical Committee. After overnight recovery in Ham’s F-10 containing 6.1 mmol/L glucose, 10% FBS, 2 mmol/L GlutaMAX, 0.75% BSA, 50 units/mL penicillin, and 50 μg/mL streptomycin, human islets were dispersed for RNA silencing. The percentage of beta cells, assessed by insulin immunofluorescence, was 50 ± 11% (mean ± SD).

#### Beta cell differentiation and immunocytochemistry

Control iPSC lines HEL115.6 (Cosentino et al. 2018) and 1023A (Lytrivi et al. 2021) were differentiated into beta cells using a previously published 7-step protocol (Fantuzzi et al. 2022; De Franco et al. 2020). At the end of the stage 4, cells were seeded into 24-well Aggrewell 400 microwell plates (Stem Cell Technologies) at a density of 0.9 ·10^6^ cells per well after which differentiation was carried out as described previously (Cosentino et al. 2018).

Stage 7 differentiated beta cells were washed twice with PBS containing 0.5 mM EDTA and incubated in Accumax (SIGMA #A7089) for 8 min at 37 °C after which 50% volume of KnockOut Serum Replacement (ThermoFisher #10828028) was added to stop the reaction. After centrifugation at 400 g for 5 min at room temperature, cells were resuspended in 1 mL HAM’s F-10 medium, supplemented as indicated before (Demine *et al*., 2020). 75,000 cells in 500 μL medium were seeded per square of a Nunc Lab-Tek II ICC chamber (ThermoFisher).

Immunohistochemistry analyses were carried out largely as described previously (Demine et al. 2020), using the following primary antibodies: INS (Dako (Agilent), IR002, Ready to use solution), GCG (Sigma-Aldrich, G2654, 1/1000) and JUND (Atlas Antibodies, HPA063029, 1/100) and secondary antibodies A-11075, A-21206, A-31571 (ThermoFisher, 1/500). Pictures were taken using an Axiovert fluorescence microscope (Zeiss).

#### RNA silencing in dispersed human islets and hiPSC-derived islets

Dispersed human islets and hiPSC-derived islets were transfected with siRNAs for human JUND, (Dharmacon, LQ-003900-00-0002) or human BHLHE41 (Dharmacon, LQ-010043-00-0002). siRNA with no homology to any known mammalian gene was used as negative control (AllStars Negative Control siRNA, Qiagen). siRNA-Lipofectamine RNAiMAX (Invitrogen, Life Technologies) complexes were formed in Opti-MEM and diluted four times in Ham’s F-10 medium without BSA or penicillin-streptomycin. Transfection was done using 30 nM siRNA and Lipofectamine at a final dilution of 1/250. Medium was refreshed after 16 hours and cells were studied after another 48 h.

#### Assessment of cell apoptosis

The percentage of apoptotic cells was determined by nuclei staining with propidium iodide (5 µg/ml in PBS, Sigma-Aldrich) and Hoechst 33342 (10 µg/ml in PBS, Sigma-Aldrich), as described (Lytrivi et al. 2021). A minimum of 500 cells were counted for each experiment by two operators.

#### mRNA extraction and real time PCR

Total RNA was extracted using the RNeasy Plus micro kit (Qiagen), according to the manufacturer’s instructions. Briefly, cells were washed once with PBS and collected in 100 µl of the kit’s lysis buffer. RNA was eluted in 14 µl and retro-transcribed as described previously (Vanheer et al. 2019). Primer sequences are listed in Table S7. All assays had an efficiency above 95%. Relative quantities of each transcript were calculated as arbitrary units from comparison to the standard curve. Relative expression level of the target transcript was presented as the ratio of the target transcript quantity to the housekeeping transcript quantity (GAPDH, ACTB).

#### Immunohistochemistry

Briefly, 4 µm tissue sections were retrieved from 4% formalin-fixed, paraffin-embedded tissue blocks of normal pancreas tissue (68-year-old male, 59 year-old-male and 64-year-old female) from the biobank of the University Hospital in Leuven (Belgium). Two pathologists (M.V.H. and T.R.) separately evaluated all histological sections. This study was approved by the ethical committee of the University Hospital in Leuven (Belgium) (S32980).

Immunohistochemistry analyses were carried out largely as described previously (Ceulemans et al. 2017), using primary antibodies against the following proteins: ASCL2 (Merck, MAB4418, clone 7E2, 1/5000), JUND (Atlas Antibodies, HPA063029, 1/50), HEYL (Atlas Antibodies, HPA076960, 1/200) and INS (Agilent, IR002, 1/100). Pictures were taken using a Leica DMLB (Leica Microsystems).

#### SCope

The integrated scRNA-seq data and pySCENIC results can be explored interactively in SCope (Davie *et al*., 2018). Loompy (v2.0.17) (Linnarsson Lab., 2015) was used to create the loom files which were uploaded to SCope. The embedding of the regulon and integrated gene expression based UMAP clustering, as seen in this article, were added to the loom file.

---

**Figure S1.**
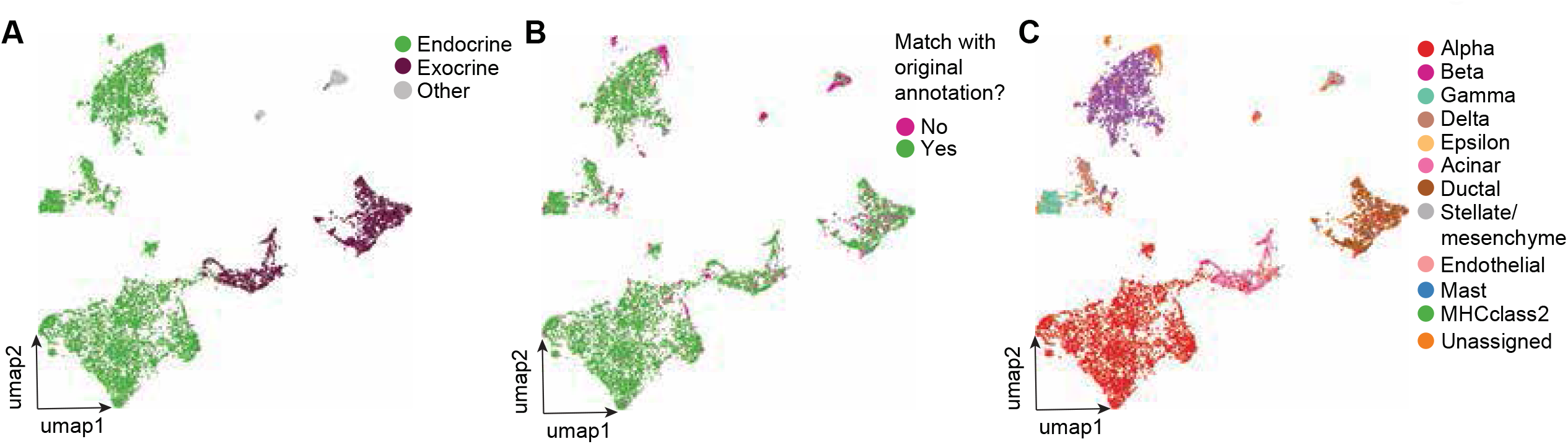
Integrated Analysis of scRNA-seq Identifies 12 Pancreatic Cell Types. **(A)** Gene expression based UMAP of 7393 single cells colored by pancreatic lineage. Endocrine cells include alpha, beta, epsilon, gamma and delta cells. Exocrine cells include acinar and ductal cells. Other cells represent schwann, MHC class 2, mast, stellate and endothelial cells. **(B)** Gene expression based UMAP of 7393 single cells showing the large degree of overlap of novel and original cell type annotation. Cells were annotated as matching when the novel cell type annotation corresponded with the original annotation from each paper. **(C)** Gene expression based UMAP of 7393 single cells annotated by the original cell type annotation defined in the individual papers.

**Figure S2.**
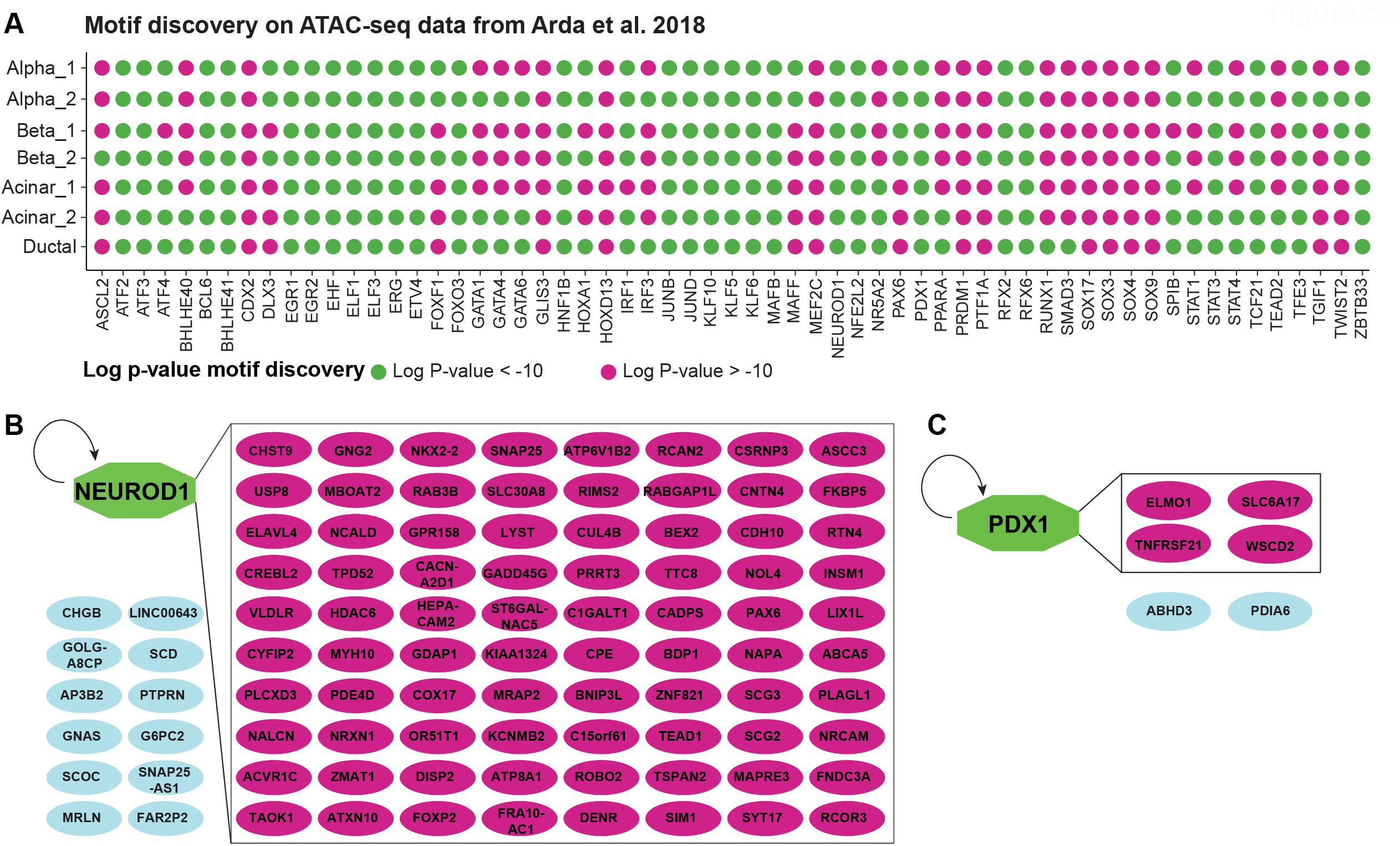
Reconstruction of Gene Regulatory Networks in the Human Pancreas Predicts a Small Set of Essential Regulators for Each Cell Type. **(A)** Bubble plot summarizing the enrichment of pySCENIC related transcription factor motifs. Bulk ATAC-seq data of FACS sorted alpha, beta, ductal and acinar cells from (Arda et al., 2018) was downloaded and analysed after which motif discovery by HOMER based motif discovery was performed on the global open chromatin landscape. Significantly enriched motifs, with logarithmic p-values below -10, are represented in green. TF motifs with logarithmic p-values above -10 were considered not significant and are represented in purple. **(B)** Extended analysis of putative NEUROD1 target genes. NEUROD1 is colored green. Target genes are color coded (purple/blue) based on whether they are part or not of the network as defined by iRegulon. **(C)** Extended analysis of putative PDX1 target genes. PDX1 is colored green. Target genes are color coded (purple/blue) based on whether they are part or not of the network as defined by iRegulon.

**Figure S3.**
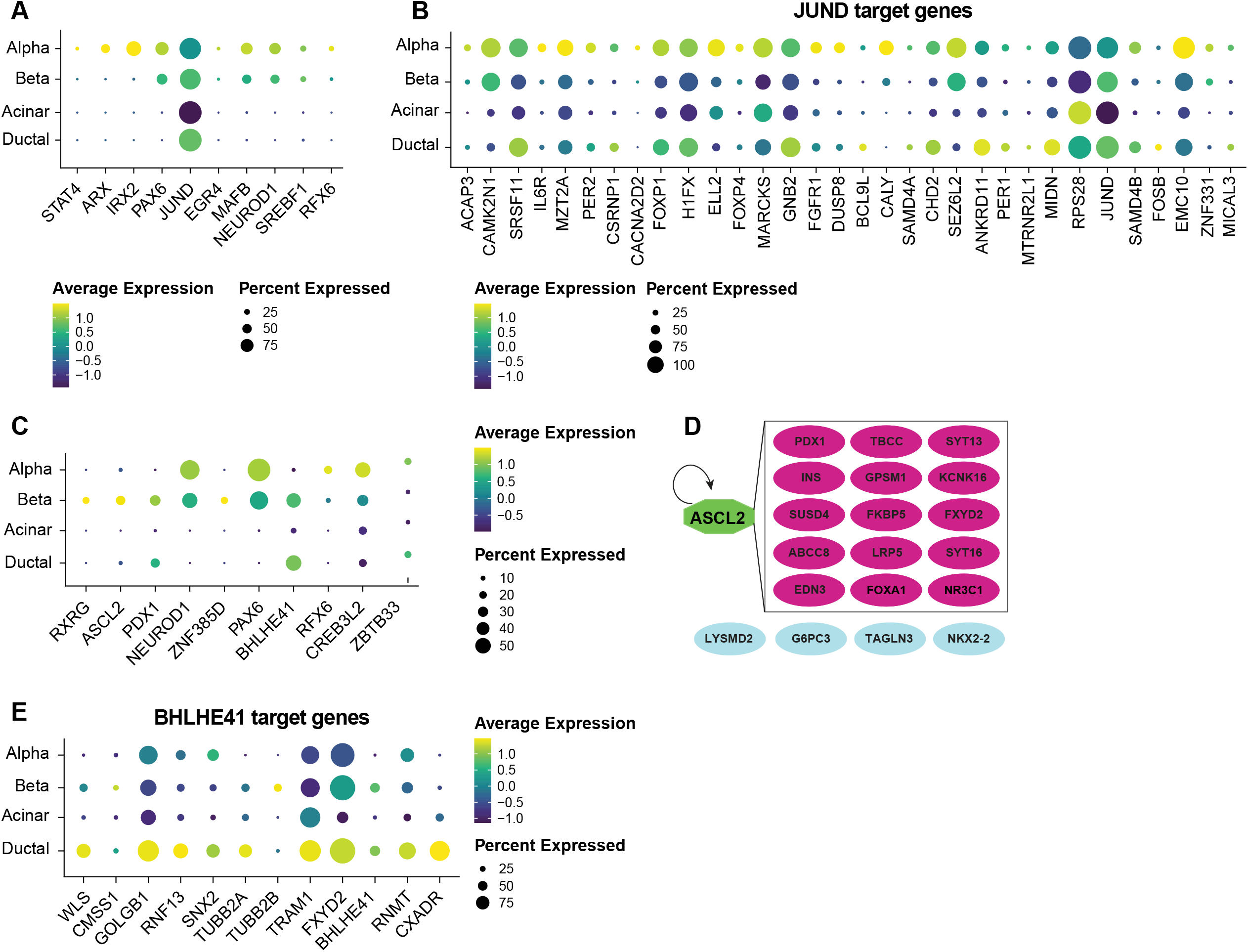
Prediction of regulators of endocrine cell identity in the human adult pancreas. **(A)** Bubble plot showing the expression of transcription factors in the HPAP dataset for the top 10 alpha cell regulons. The bubble size is proportional to the percentage of cells that express a specific transcription factor with the color scale representing the average gene expression within the specific cell population. **(B)** Bubble plot showing the expression of JUND target genes in the HPAP dataset. The bubble size is proportional to the percentage of cells that express a specific target gene with the color scale representing the average gene expression within the specific cell population. **(C)** Bubble plot showing the expression of transcription factors in the HPAP dataset for the top 10 beta cell regulons. The bubble size is proportional to the percentage of cells that express a specific transcription factor with the color scale representing the average gene expression within the specific cell population. **(D)** Extended analysis of the target genes of the ASCL2 regulon by iRegulon. ASCL2 is colored green. Target genes are color coded (purple/blue) based on whether they are part of a network. **(E)** Bubble plot showing the expression of BHLHE41 target genes in HPAP dataset. The bubble size is proportional to the percentage of cells that express a specific target gene with the color scale representing the average gene expression within the specific cell population.

**Figure S4.**
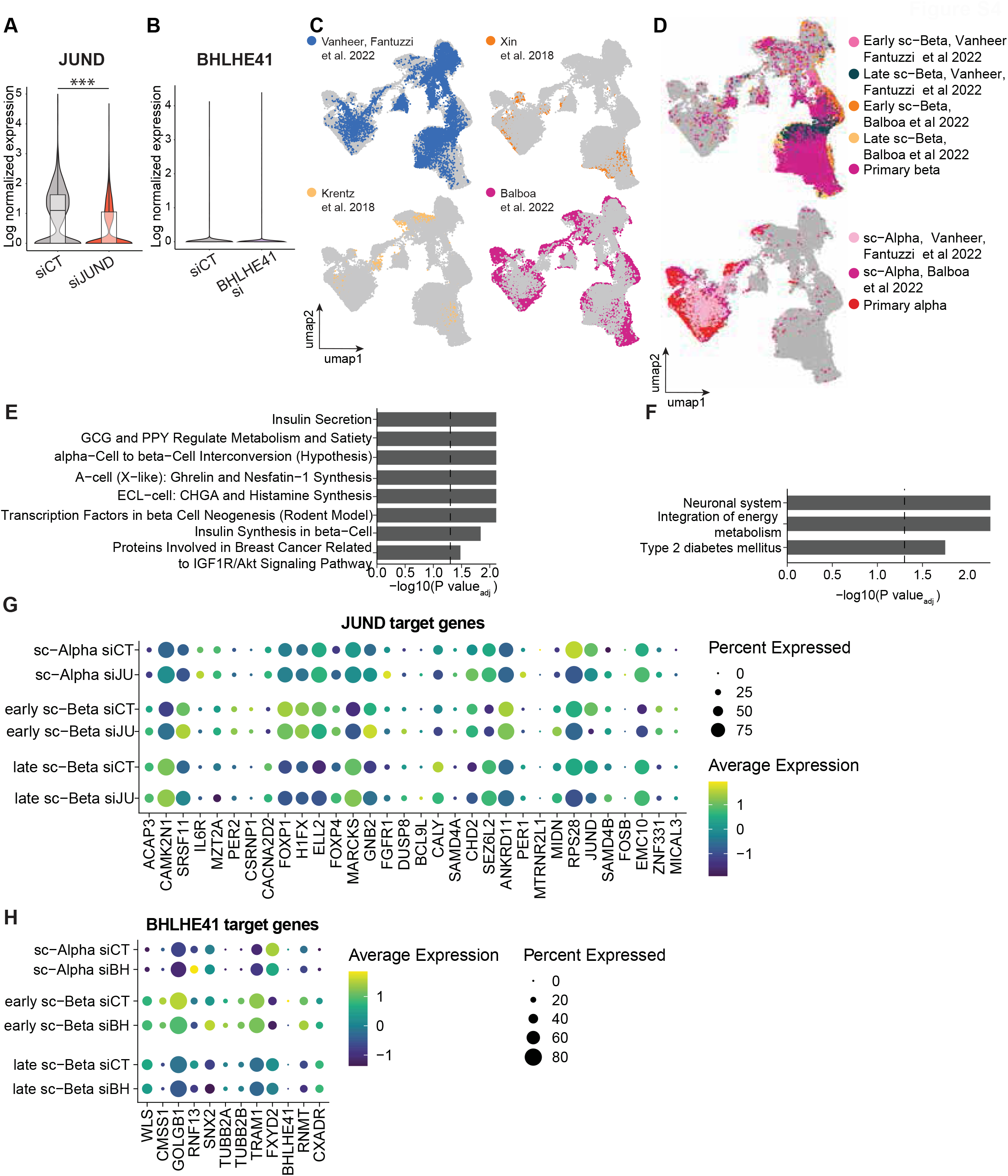
BHLHE41 and JUND depletion in hiPSC-derived islet cells. **(A-B)** Violin plot of JUND **(A)** and BHLHE41 **(B)** expression by scRNA-seq, indicating a significant decrease in JUND in siJUND transfected cells. **(C-D)** Integrated gene expression based UMAP of hiPSC islet data of Balboa et al. 2022 with this study annotated by dataset of origin **(C)** or cell type **(D)**. **(E)** GO enrichment analysis of differentially expressed genes that are upregulated in siJUND transfected cells compared to siCT control. **(F)** GO enrichment analysis of differentially expressed genes that are upregulated in siBHLHE41 transfected cells compared to siCT control. **(G-H)** Bubble plot showing the expression of JUND **(G)** and BHLHE41 **(H)** regulon target genes as predicted by pySCENIC. The bubble size is proportional to the percentage of cells that express a specific target gene with the color scale representing the average scaled gene expression within the specific cell population.

**Figure S5.**
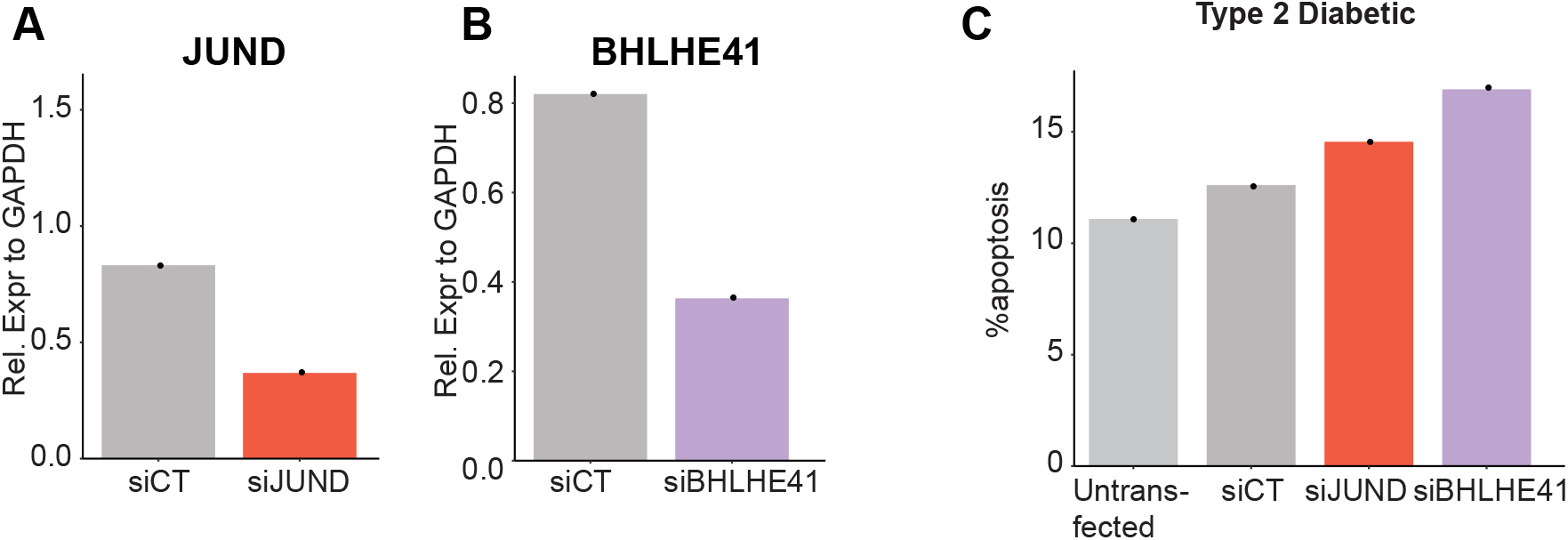
JUND and BHLHE41 mediated knockdown in type 2 diabetic primary pancreatic islets. **(A-B)** JUND **(A)** and BHLHE41 **(B)** transcript level 72 hours in human islets from a type 2 diabetic donor after siRNA transfection by RT-qPCR. **(C)** Percentage of apoptotic cells 72 hours after siRNA transfection. n = 1 islet preparation.

**Figure S6.**
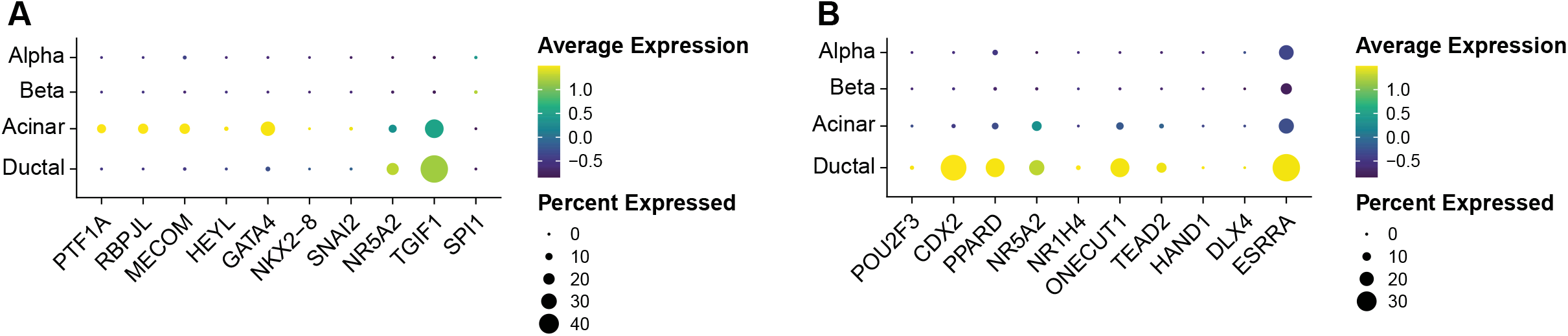
Validation of exocrine cell regulons in HPAP dataset. **(A)** Expression of transcription factors in the HPAP dataset for the top 10 acinar regulons. The bubble size is proportional to the percentage of cells that express a specific transcription factor with the color scale representing the average gene expression within the specific cell population. **(B)** Expression of transcription factors in the HPAP dataset for the top 10 ductal regulons. The bubble size is proportional to the percentage of cells that express a specific transcription factor with the color scale representing the average gene expression within the specific cell population.

## Supplementary files and tables

**Table S1. Overview of donor information and number of cell types identified for each donor**.

This table is related to Figure 1, 2, 3 and 6.

**Table S2. Overview of marker genes used to annotate cell populations**.

This table is related to Figure 1.

**Table S3. Overview of all 142 regulons**.

This table is related to Figure 2, 3 and 6.

**Table S4. Overview of regulon ranking**.

List of regulon specificity scores for all 142 regulons of non-diabetic alpha, beta, acinar and ductal cells.

This table is related to Figure 3 and 6.

**Table S5. List of putative target genes for each regulon**.

List of putative target č genes for each regulon. This table is related to Figure 2, 3, 4 and 5.

**Table S6: Percentage of cell types identified by scRNA-seq of siRNA transfected hiPSC-derived islets**.

Table with cell type percentages of siRNA transfected hiPSC-derived islets per siRNA condition. This table is related to Figure 4G.

**Table S7. Primer sequences**.

